# Age-dependent effects of infection on survival of a wild rodent reservoir host

**DOI:** 10.64898/2026.03.17.712390

**Authors:** Katherine E. Wearing, Jasmine S. M. Veitch, Janine Mistrick, Dyess F. Harp, Brendan B. Haile, Christina G. Fragel, Tarja Sironen, Meggan E. Craft, Clayton E. Cressler, Richard J. Hall, Sarah A. Budischak, Kristian M. Forbes

## Abstract

Due to long co-evolutionary histories, many zoonotic pathogens are thought to exert little or no negative effects on their wildlife reservoir hosts. However, there remains a lack of rigorous investigations in natural settings. We conducted a 3-year factorial field experiment to investigate how survival of the Puumala hantavirus (PUUV) reservoir, the bank vole, is impacted by PUUV infection, nematode infections, and food availability. We hypothesized that PUUV would not impact survival, but that coinfection with nematodes would negatively impact survival, and that increased food availability would mitigate the negative effects of coinfection. Surprisingly, we demonstrated that PUUV infected voles had substantially reduced survival when compared to uninfected voles, and this strong negative effect manifested in young voles. Nematode removal increased survival of young voles and food supplementation interacted with movement rather than survival. Our results provide empirical evidence in a natural system for infection reducing survival of its reservoir host.

## Introduction

Due to their long co-evolutionary history, many zoonotic pathogens are thought to be largely asymptomatic to their reservoir hosts. This pattern has played out across many high-profile zoonotic systems, such as Hendra virus, Nipah virus, Sin Nombre virus, Puumala hantavirus, Hantaan virus, tick-borne encephalitis virus, and Lassa virus [1–8]. However, much of this research is based on experiments (often with very small sample sizes) conducted in a laboratory, which lack natural stressors. There are few studies in field settings that address the potential negative impacts of pathogens on their reservoir hosts (but see [9, 10]).

Powerful field studies, with spatial and temporal replication and potentially experimental manipulation, are essential to discern the effects of factors influencing reservoir hosts. Large data sets and long-term sampling in wild settings can overcome noisy data that may obscure subtle differences between hosts with different characteristics or occupying different environments. However, these types of long-term studies examining survival of reservoir hosts, and especially incorporating experimental manipulation, are rare in the scientific literature.

Many hantaviruses share long coevolutionary histories with their rodent hosts, and are considered textbook examples of pathogens that cause asymptomatic infections in their reservoir hosts [11]. For example, Puumala hantavirus (PUUV) is a zoonotic virus which shares a long coevolutionary history with its bank vole (*Clethrionomys glareolus*) reservoir host [12]. While PUUV causes acute disease in humans, it causes chronic and apparently asymptomatic infections in the bank vole reservoir ([13] but see [9]). The highest reported global incidence of PUUV-caused human disease is in Finland, where 1,000 to 3,000 cases are reported annually [14]. This reflects high PUUV prevalence in voles, as every human case is an individual spillover event [13], and emphasizes the public health importance of understanding infection biology in the reservoir host.

There is a lack of experimental research on how ecological factors may modulate the effects of infections on reservoirs and other host types. Two key factors which could affect survival of reservoir hosts are food and coinfections. Nematodes and viruses activate conflicting branches of the vertebrate immune system [15], so coinfections may have different outcomes from single infections. Bank voles can be naturally coinfected with nematodes and PUUV [16]. Given that immune activation is energetically costly [17], and that food may change a host’s ability to tolerate infection [18], food availability may modulate relationships between infections and host survival. Indeed, supplemental food has been shown to increase immune responses [19] and increase pathogen resistance [20] to nematode infections in wild rodents.

To address the lack of research on reservoir host survival in natural environments, we conducted a 3-year field experiment in Finland, investigating how PUUV infection, nematode coinfection, and food availability impact bank vole survival. We experimentally removed nematodes and provided supplemental food to wild bank vole populations in a powerful two-factor study design. Given existing evidence from laboratory studies, we hypothesized that PUUV infection would not alter bank vole survival. However, we predicted that coinfections with nematodes would negatively impact survival and that food supplementation would mitigate the negative effects of coinfections.

## Methods

### Study system and setting

Bank voles are a common small mammal in forests of northern Europe and are naturally infected with PUUV and nematodes. Individuals usually live for less than a year [21] and their population abundance fluctuates seasonally. Reproduction begins in spring [22], with population sizes peaking in late summer or fall and then declining dramatically during the winter period. Reproductively active voles maintain territories, and other voles may disperse from their natal habitat, although dispersal depends on quality of natal habitat [23]. Voles become infected with PUUV after inhaling aerosolized virus shed by infected voles in their urine, saliva and feces [13]. Nematodes are directly transmitted between bank voles when they contact feces containing nematode eggs/larvae or through contact between infected voles [24].

Our experiment was conducted in 12 forest sites in southern Finland (61.0775°□ N, 25.0110°□E, based out of the Lammi Biological Station). Study sites were located in mature forest patches at least 2-hectares in size and at least 2-kilometers apart to prevent vole movement between sites. At each site, we established a 100-meter x 100-meter trapping grid to administer treatments and monitor voles. Ugglan multi-capture live traps (Grahnab, Gnosjö, Sweden) were placed in a permanent configuration with 61 traps evenly spaced (Supplemental Figure 1). Our trapping grids were not fenced and allowed for the natural movement of voles and other animals in the environment.

### Experiment design and longitudinal monitoring

We conducted a fully factorial field experiment manipulating food availability and coinfection. We manipulated food availability by adding supplemental food to the environment (‘fed treatment’), and coinfection by removing nematodes from voles (‘deworm treatment’). Altogether, we had four treatment groups: 1) fed/deworm, 2) fed/control, 3) unfed/deworm, 4) unfed/control (for additional details see [25]), with three replicate populations of each treatment group. The 12 vole populations/sites were randomized to treatment groups. Two sites were replaced with new locations during the experiment due to logging and unusually low vole numbers, giving us 14 sites overall, but 12 sites at any one time point. One site was logged prior to our final visit in May 2024, resulting in only 11 sites trapped for that month.

The experiment was conducted from May 2021 through May 2024. Voles were longitudinally monitored at monthly intervals from May through October each year. At other times of year, snow cover in the forests prevented access to the field sites.

We used a capture-mark-recapture approach, such that at first capture, voles received a subcutaneous Passive Integrated Transponder (PIT tag; ‘Skinny’ PIT Tag, Oregon RFID, Milwaukie, USA) for future identification. Within a monthly trapping occasion, traps were set in the evening, then checked and reset in the morning and evening for the next two days (four checks total per trapping occasion). Upon capture, we scanned voles for a PIT tag and recorded the following information: 1) PIT tag number, 2) sex, 3) reproductive status, 3) head width, and 4) vole weight. Additionally, we collected blood samples and saliva samples to test for PUUV infection and virus shedding, respectively, and fecal samples to screen for nematode eggs. Following processing, voles were released at their point of capture. If a vole was captured again that month, we recorded its presence and released it without further sampling. Traps were replaced and cleaned following each capture to prevent contamination of fecal samples.

To remove nematodes from voles at deworm sites, we administered a combination of Ivermectin (10mg/kg; [26]) and Pyrantel (100mg/kg; [27]), which treats both larval-and adult-stage infections [20]. In a separate analysis, Veitch et al [28 Preprint] demonstrated that this deworm treatment successfully reduced the likelihood of a vole being infected with nematodes, decreased the prevalence of nematodes in the population, and reduced the intensity of nematode infections among infected voles. Since Ivermectin can also treat tick infections, we ran a validation analysis which indicated no relationship between deworm treatment and tick infection (see Supplement for details), allowing us to attribute the effects of our deworming medication to nematode infections. At control sites, voles were given a placebo treatment of sugar water instead of deworm.

Supplemental food was provided at fed sites once every two weeks from May through the middle of November of each year. For this, we scattered 7.875kg of mouse chow pellets (3227 kcal/kg; Altromin Spezialfutter GmbH & Co. KG, Lage, Germany) and 7.875kg sunflower seeds (6350 kcal/kg) evenly across the entire grid (as described in [25] and similar to [20]). The sunflower seeds ensured that supplemental food was still available if rains occurred soon after distribution.

### Pathogen diagnostics

We determined hantavirus infection using immunofluorescence assays (IFA). Since bank voles are chronically infected with PUUV, antibody detection is effective for determining current PUUV infection [29,30]. IFAs were run on vole blood serum based on validated protocols [31,32]. Following identification of seropositive voles, we conducted an analysis to distinguish those with maternal antibodies from actual infection. We used a generalized additive mixed model to determine the mass below which voles might have maternal antibodies (determined to be 13.8g, Supplemental Figure 2), following methods in Voutilainen et al [33]. We validated our main analyses by removing any PUUV seropositive (and PUUV RNA negative) samples from voles weighing <13.8g and rerunning the analyses to confirm that potential maternal antibodies were not driving the results (Supplemental Tables 1, 2, 3, 4). A small proportion of voles (32/∼1500 individuals, ∼2%) showed an unexpected pattern of seropositivity (switched from positive to negative) and were removed from all analyses independent of mass (See Supplement for more details). Additionally, there were 12 captures out of ∼2200 where we could not test a vole for PUUV infection, but in subsequent months the same vole tested negative for infection. In these 12 cases, voles were designated as negative since PUUV causes chronic infections.

Nematode infection status and intensity were determined through fecal egg counts (FECs) using a salt flotation method ([34], similar to [35,36]). Infection intensity was measured in eggs per gram of feces, and fecal samples weighing less than 0.01-grams were removed from all analyses as they have been shown to be less accurate for measuring nematode infection status and intensity [28 Preprint]. Separate analyses in this study system also showed that nematode fecal egg count is correlated with adult worm count in these populations (Budischak, unpublished data).

### Data analyses

We used Cox proportional hazards models (R package “survival” [37]) to investigate vole survival. These models can provide mortality hazard ratios (a proxy for survival) which describe the chance of an individual dying at a given time, if they have already survived to that time point, in relation to the model covariates. We assigned voles as “dead” if they disappeared the next month and were never captured again. As such, vole “survival” and “mortality” discussed in this paper specifically references apparent survival and apparent mortality, as is standard for wild small mammals.

Since PUUV infection status could change over the course of the study (negative to positive), we fit Cox proportional hazards models with time-varying covariates, following methods presented in Zhang et al [38]. In these models, the hazard of an event (i.e. apparent death) is measured for all individuals at each time interval, which ends with the event either occurring or not occurring. If the event does not occur until after the regular sampling (i.e. a vole disappeared over the winter when we were not sampling monthly or after the completion of data collection in May 2024), then there is not enough information to know when an animal “died”. This situation is “right censored” and is handled by the time-varying covariate model, as it measures interval to interval. In our models, data were clustered by individual vole ID, allowing risk error calculations to account for multiple sampling of the same individuals. We checked the proportional hazards assumption for the models and added time transformations as needed (allowing covariates which violated the proportional hazards assumption to have relationships that changed over time).

Since voles that did not survive long enough to be captured were automatically excluded from the data set (an example of left truncation) we needed to estimate vole age at first capture to avoid biasing our models toward individuals with longer survival times. We used the age-head width relationship described in Kallio et al [39] to fit a generalized additive model with vole age as the response and vole head width as the explanatory variable. Predictions from this model were used to estimate vole age based on the corrected head width measure at a vole’s first capture (see Supplement for more information). Rounded vole age (in months) at first capture was used to estimate age at subsequent captures and was included as the starting time in the Cox proportional hazards models.

With the Cox proportional hazards models, we ran a primary analysis on the full data, a secondary analysis with subsetted data and several validation analyses. Additionally, we ran an analysis with generalized linear mixed models to investigate vole overwinter survival. For each of these analyses (primary, secondary, validation and overwinter), we expected that PUUV infection, deworm treatment, food treatment, vole sex, nematode infection, reproductive status and month could all relate to vole survival, so these covariates were included in each model. We fit a set of 46 candidate models with different combinations of interactions (Supplemental Table 5) and compared them using corrected Akaike’s Information Criterion (AICc). Based on our experimental manipulations, focus on infection, and knowledge of vole biology, we were specifically interested in interactions involving deworm treatment, food treatment, PUUV infection, nematode infection and vole sex. So, we compared all combinations of 2- and 3-way interactions between these factors (Supplemental Table 5). Inference was made based on the model favored by AICc which had the highest AICc weight. To help interpret interaction terms, we plotted predicted relative proportional hazard ratios from the favored model using “ggplot” [40] and made pairwise comparisons of estimated marginal means from the Cox proportional hazards models using “pairs” and “emmeans” [41]. A Bonferroni correction was included in post hoc pairwise comparisons to adjust for multiple comparisons.

In the secondary analysis, we examined mortality hazard among younger and older voles. A previous study with a sub-set of these data (with PUUV infected voles removed) indicated that the relationship between nematode infection and the likelihood that a vole would be captured again may change with vole age [28 Preprint]. The mean apparent lifespan of voles in our study was between 5 and 6 months, and there was a 7-month interval between October trapping and May trapping, so a vole that was 7 months old had lived longer than average and had often survived an overwinter period. So, we divided the data into younger voles (less than 7 months old, 78% of the observations), and older voles (7 or more months old, 22% of the observations).

We ran validation analyses to check whether the results were driven by vole movement rather than true mortality. In particular, we investigated the potential role of transient adult voles, which we defined as reproductively active voles that were only captured once. Since voles normally maintain territories when they are reproductively active [42], being captured only once could suggest that animals which moved onto sites only briefly (e.g. to access supplemental food) were then leaving, rather than dying. To address this, we ran the primary and secondary analyses again with these transient voles removed.

We also investigated overwinter survival. Following similar methods to Kallio et al [9], our response variable was whether or not voles caught in October were caught again the following May. Measures of PUUV infection and nematode infection in October, as well as experimental treatments and vole sex were included as predictors. A set of 46 candidate generalized linear mixed models were run with the same interactions as the primary analysis (3 of 46 failed to converge and were excluded from model comparison to avoid biasing the AICc; Supplemental Table 5). The candidate models had binomial distributions and study site included as a random effect (fit with R package ‘glmmTMB’ [43]). Models were compared using AICc.

## Results

### Summary Results

During the study, we caught 1489 individual voles. Of these individuals, 33% were recaptured at least once and we had 2198 total captures. Twenty percent (20%) of the total captures were positive for PUUV while 32% of the total captures were positive for nematode infections. Eight percent (8%) of the total captures were coinfected with PUUV and nematodes.

We determined month of apparent death for 849 voles. Of these, 27% had died by the time they were 2 months old, 43% had died by the time they were 3 months old, and 68% had died by the time they were 6 months old (see Supplemental Figure 3 for a visualization of how many voles died in each month). Thirty-two percent (32%) of the 849 voles survived for 7 months or longer.

### Infection Results

PUUV infection increased the mortality hazard, and this trend was driven by younger voles (less than 7 months old). In the primary analysis, the AICc favored model with the highest AICc weight did not include any interactions between covariates (M1, Supplemental Table 6). In this model, the chance of a PUUV infected vole dying at a given time (if they had survived to that time point) was 59% higher than a PUUV uninfected vole (hazard ratio=1.59, p<0.0001, Supplemental Table 7). However, this result was driven by the effect of PUUV infection on younger voles. For younger voles (secondary analysis), the favored model with the highest AICc weight included a nonsignificant interaction between PUUV infection and vole sex (MY5, Supplemental Table 8, Supplemental Table 9). Post hoc pairwise comparisons with the non-significant interaction revealed that every PUUV infected group had significantly higher mortality hazard than every PUUV uninfected group, regardless of vole sex (Supplemental Figure 4). An AICc favored model with slightly lower AICc weight, which did not include an interaction with PUUV, was used to plot the relationship between PUUV infection and younger vole mortality hazard in order to avoid plotting main effects of PUUV when it was part of a nonsignificant interaction (Figure 1). This model confirmed that younger voles infected with PUUV had a much higher mortality hazard than uninfected younger voles (MY1, hazard ratio=1.73, p<0.0001, Figure 1). In contrast, PUUV infection was not related to mortality hazard in older voles. Among older voles, there were two AICc favored models which shared the highest AICc weight, MO9 and MO21. Neither of these models included an interaction involving PUUV infection status (Supplemental Table 10) and in both models, there was no relationship between PUUV infection status and mortality hazard (MO9, p=0.36; MO21, p=0.31; Figure 1, Supplemental Table 9, Supplemental Table 11). In the overwinter survival analysis, no model was favored over the null model (AICc) indicating that none of the predictors helped to explain the variance in likelihood of surviving the winter period (Supplemental Table 12).

**Figure 1.**
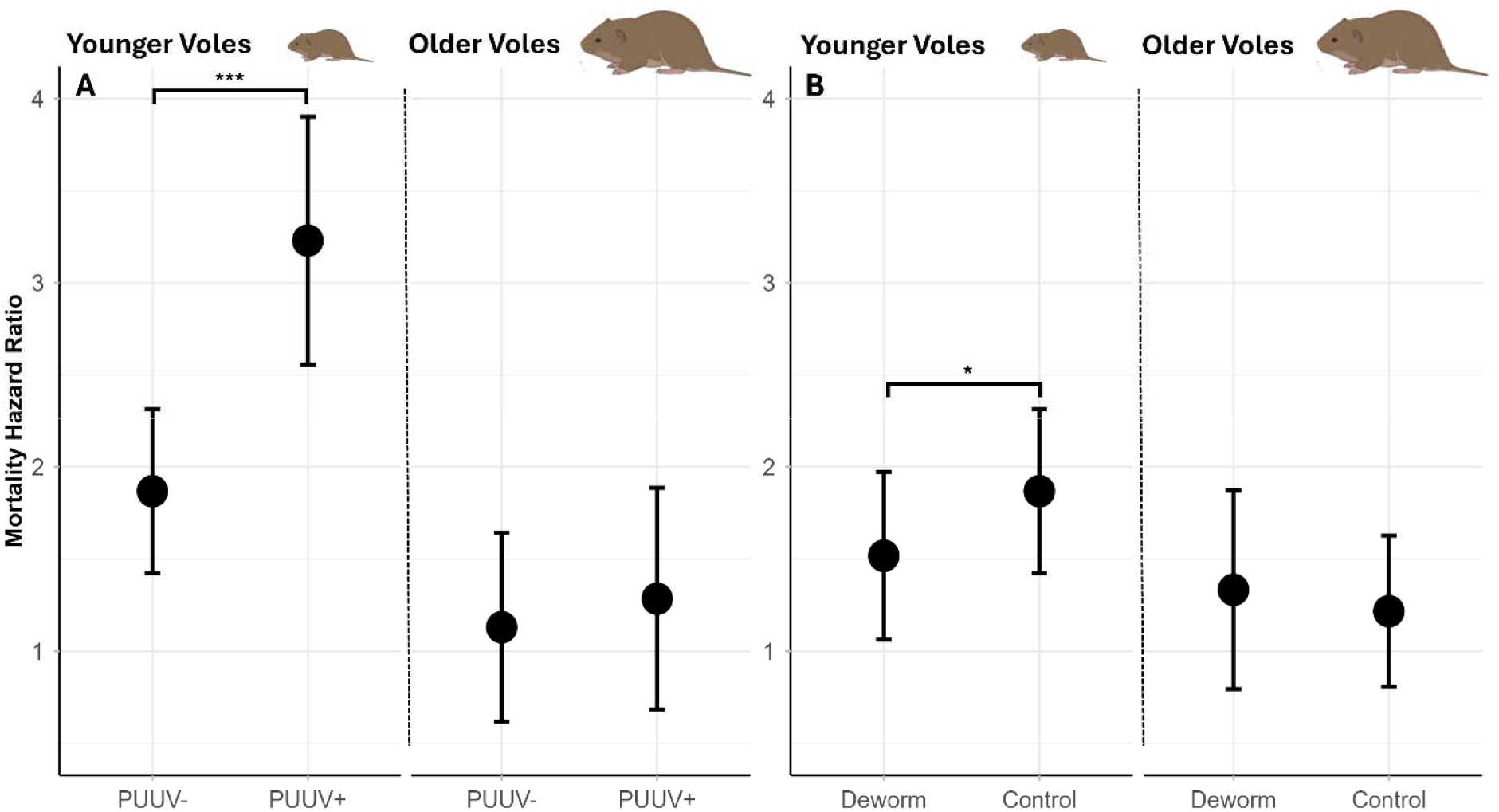
Relationships between PUUV infection and chance of dying at a given time (Panel A), and between deworm treatment and chance of dying at a given time (Panel B), are different for younger voles (<7 months old) and older voles (7 or more months old). This plot shows predicted mortality hazard ratios, relative to the sample average for all predictors in the Cox proportional hazards models (MY1 and MO9). The relative mortality hazard is higher among younger PUUV infected voles (p<0.0001) while we detected no difference in mortality hazard with PUUV infection in older voles (p=0.36). With deworm treatment (Panel B), the relative mortality hazard is lower in younger voles (p<0.05) while we detected no difference in mortality hazard with deworm treatment in older voles (p=0.48). Error bars on both panels indicate 95% confidence intervals.

Deworm treatment decreased the mortality hazard among younger voles. Although deworm treatment did not significantly impact the mortality hazard in the primary analysis (M1, p=0.085; Supplemental Table 7), the secondary analysis revealed that the relationship between deworm treatment and mortality hazard changed with vole age. Younger voles at deworm sites had a 19% lower chance of dying at a given time (if they had already survived to that time) than younger voles at control sites (MY5, hazard ratio=0.81, p<0.05; Figure 1, Supplemental Table 9). Although one of the favored models for older voles contained a nonsignificant interaction between deworm treatment and nematode infection status, post hoc pairwise comparison showed no significant differences between any groups (MO21). Additionally, the other favored model for older voles, which had no interactions involving deworm treatment, confirmed that the treatment did not impact the mortality hazard among older voles (MO9, p=0.48; Figure 1). Surprisingly, younger voles infected with nematodes also had a lower mortality hazard (MY5, hazard ratio=0.82, p<0.05; Supplemental Table 9). Nematode infection status did not have a significant relationship with mortality hazard among older voles (MO9, p=0.59, Supplemental Table 9).

### Vole Characteristics and Environment

Mortality hazard was also related to vole characteristics and season, independent of PUUV infection status. In the primary analysis, male voles had a significantly higher mortality hazard than female voles (M1, hazard ratio=1.22, p<0.01, Figure 2, Supplemental Table 7). The secondary analysis showed that the relationship between vole sex and mortality hazard may depend on other conditions in younger and older voles, as sex was included in nonsignificant interaction terms for both age groups (MY5, sex*PUUV infection, Supplemental Table 9, Supplemental Figure 4; MO9, sex*food treatment, Supplemental Table 9, Supplemental Figure 5; MO21, sex*food treatment, Supplemental Table 11, Supplemental Figure 6). The primary analysis showed that the relationship between reproductive status and mortality hazard changed over time, as the proportional hazards assumption was violated for reproductive status, requiring a time varying function. This time varying function revealed that the relationship changed when voles were 4.3 months old. When voles were less than 4.3 months old, being reproductively active decreased the mortality hazard (M1, hazard ratio=0.28, p<0.0001; Supplemental Figure 7, Supplemental Table 7). Once they were older, being reproductively active increased the mortality hazard (M1, hazard ratio=1.34, p<0.0001; Supplemental Figure 7, Supplemental Table 7). The relationship between mortality hazard and month also changed with vole age, as month violated the proportional hazards assumption. The time varying function showed that this relationship changed when voles were 12.4 months old. Voles less than 12.4 months old had a lower mortality hazard as the summer progressed (i.e., approaching October; M1, hazard ratio=0.46, p<0.0001; Supplemental Table 7). Meanwhile, voles older than 12 months had a higher mortality hazard as the summer progressed (M1, hazard ratio=1.06, p<0.0001; Supplemental Table 7), although this age group included few individuals (n=34).

**Figure 2.**
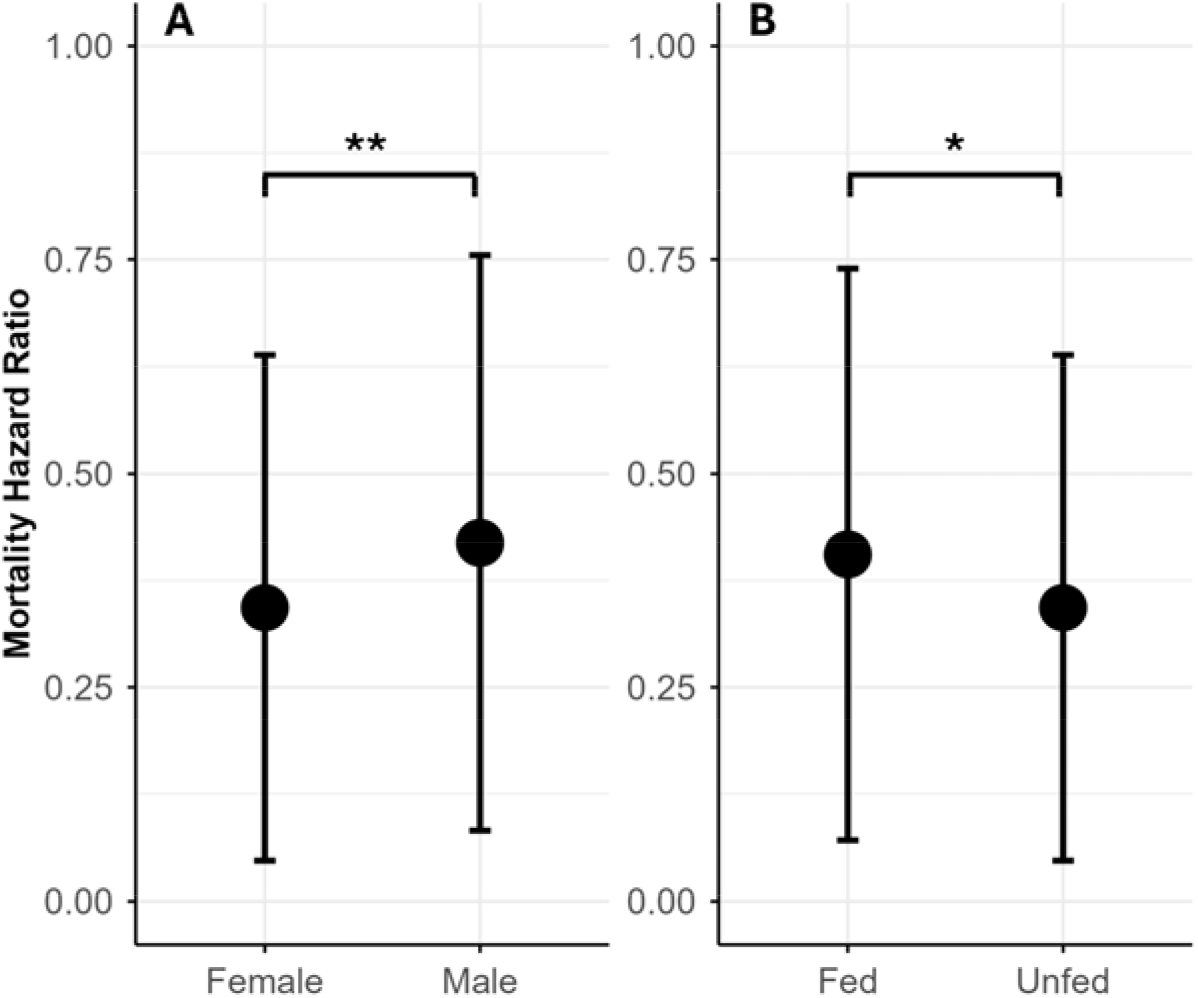
Relationships between vole characteristics/environment and mortality hazard. Panels A and B show predicted mortality hazard ratios, relative to the sample average for all predictors in the Cox proportional hazards model (M1). Panel A shows the relative mortality hazard ratios for vole sex. Panel B shows the relative mortality hazard ratios for food supplementation. Error bars indicate 95% confidence intervals for both panels.

Food supplementation increased vole mortality hazard. The primary analysis showed that voles at fed sites had an 18% higher chance of dying at a given time (if they had already survived to that time) than voles at unfed sites (with deworm treatment held at control baseline) (M1, hazard ratio=1.18, p<0.05; Figure 2, Supplemental Table 7). Food supplementation increased vole mortality hazard among younger voles (MY5, hazard ratio=1.27, p<0.01, Supplemental Table 9), but among older voles, the relationship between food treatment and mortality hazard depended on vole sex, and was a weaker relationship. Both of the highest AICc weight models for older voles included nonsignificant interactions between food treatment and vole sex (Supplemental Table 10). Post hoc pairwise comparisons with each of these models showed that the only difference between older voles at fed and unfed sites was that male voles at unfed sites had a higher mortality hazard than female voles at fed sites (Supplemental Figure 5, Supplemental Figure 6).

### Validation Analysis

With transient adult voles removed from the dataset, the relationship between PUUV infection and mortality was the same as in the main analyses, but food treatment was no longer a significant predictor of mortality hazard. This provides insight into potential effects of food treatment on vole movement versus true mortality. The favored model in the primary analysis with transient adult voles removed, T7, included a significant interaction between food treatment and deworm treatment (Supplemental Table 13, Supplemental Table 14). Deworm treatment decreased the mortality hazard only when food was not added to the environment (T7, Figure 3). Notably, there was no significant difference between the mortality hazard of voles from fed and unfed sites when they did not receive deworm treatment (T7, Figure 3). In the secondary analysis with transient adult voles removed, there was a similar interaction among younger voles, although it also depended on vole sex (TY34, food treatment*deworm treatment*sex, Supplemental Table 15, Supplemental Figure 8). The general pattern of results among older voles did not change with transient adult voles removed (Supplemental Table 16, Supplemental Table 17, Supplemental Figure 9). Importantly, in both the primary and secondary analyses with transient adult voles removed, the general relationship between PUUV infection and mortality hazard was the same as in the full primary and secondary analyses (Supplemental Tables 13, 14, 15, 16, 17). These results indicate that the PUUV-mortality relationship is not driven by transient adult voles.

**Figure 3.**
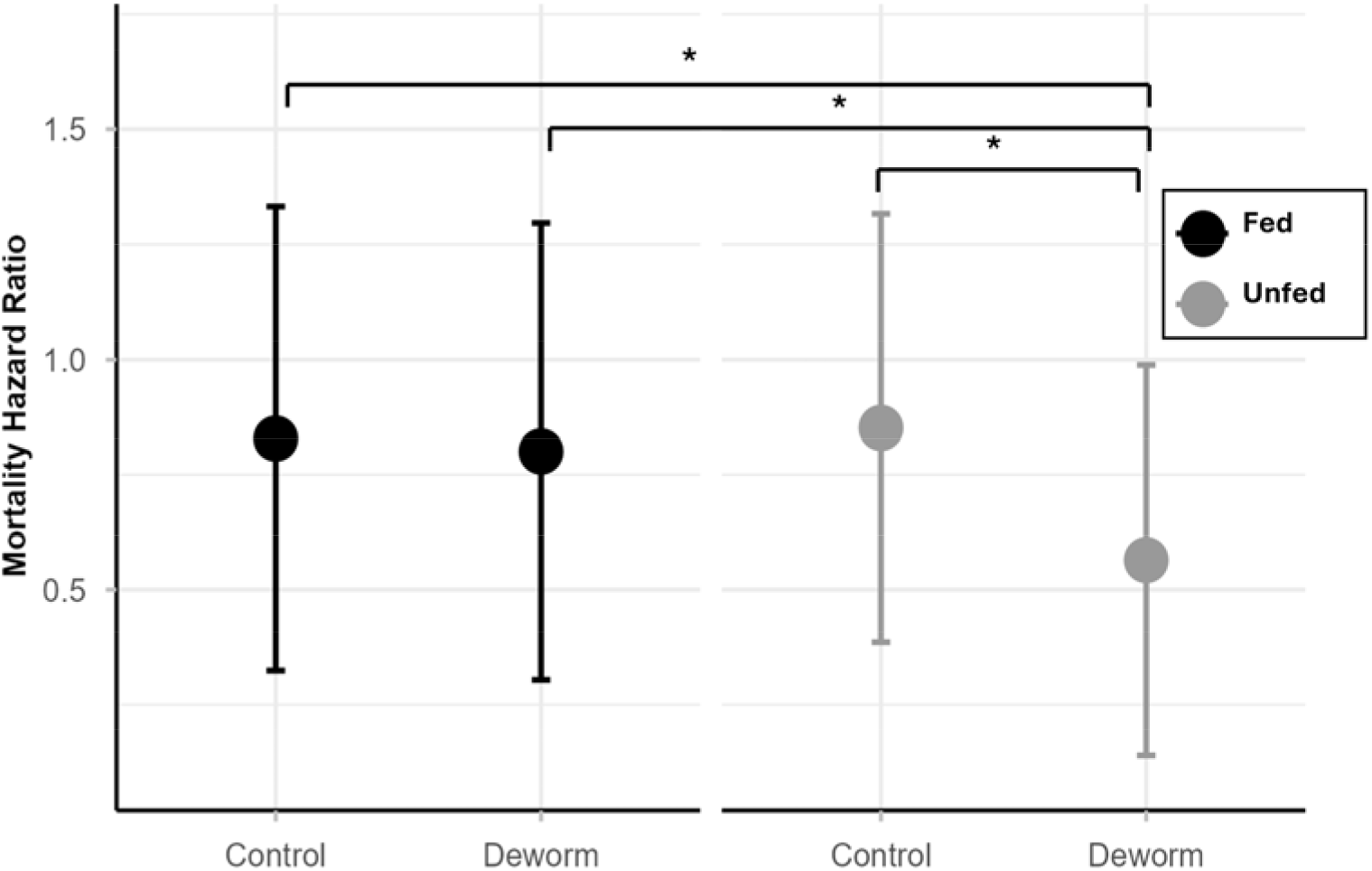
Interaction between food treatment and deworm treatment in the primary analysis with transient adult voles removed. This plot shows predicted mortality hazard ratios, relative to the sample average for all predictors in the Cox proportional hazards model (T7). Notably, we detected no difference between fed and unfed sites not receiving deworm.

## Discussion

There is a need for large-scale field studies in wild populations to characterize the potential negative effects of infection on reservoir hosts, and how these effects may be augmented by other variables, such as food availability and coinfections. We used a powerful field experiment to investigate the impact of food availability and nematode coinfections on survival of bank voles, the reservoir host for zoonotic PUUV. Although we hypothesized that PUUV infection would not decrease host survival, our results show that infected voles indeed had substantially reduced survival when compared to uninfected voles, and that this negative relationship was dependent on vole age.

Our results illustrate that PUUV reduces survival in its reservoir host, that this reduction in survival cannot be explained by harsh winter conditions or vole movement. In many systems with long co-evolutionary relationships like PUUV, reservoir hosts do not appear to be negatively affected by their pathogens, and it is humans that suffer severe negative effects following spillover infections [44]. Indeed, histopathological investigations have found that PUUV infection does not cause any detectable damage to bank voles [45]. However, Kallio et al [9] showed that PUUV reduced overwinter survival of voles in Finland. Given the harsh winter conditions in Finland, it was not clear if this reduction in survival was only apparent in winter when voles suffer severe seasonal population declines, and studies at other times of year presented conflicting results [29,46,47]. Here, we used a rigorous field study to demonstrate that PUUV can reduce bank vole survival throughout the entire year and in seemingly favorable environments. Since our grids allowed for natural vole movement, it was important to account for the potential impacts of this movement on apparent survival. We ran a validation analysis showing that transient movement of adult voles onto the grids did not impact the relationship between PUUV infection and apparent survival. This is consistent with previous studies which have found no relationship between PUUV infection and vole movement [47,48], indicating that this result reflects true mortality and not vole movement.

Nematode removal also slightly increased vole survival, but we found no evidence of an amplified coinfection effect. Very few studies have manipulated infection to examine coinfection and survival in wild populations. The one important exception is a field experiment by Ezenwa and Jolles [49], which found that nematode removal improved the survival of African buffalo infected with bovine tuberculosis. Our results show that this coinfection effect may not be a general trend. Interestingly, deworm treatment and nematode infection, which are expected to act in opposite directions, both improved survival among younger voles. A potential explanation is that nematode infections limit dispersal of young voles, thus “improving survival”. This counterintuitive result highlights the importance of experimental manipulation when discerning pathogen effects on survival; older voles are more likely to be infected with nematodes [28 Preprint] and older individuals have higher survival, creating a (non-causal) correlation between survival and nematode infection.

A key finding of our study is that the negative effects of infections only occurred in certain age cohorts. There is a lack of research on changes in infection survival with age in wildlife, particularly mammals. A small number of studies have reported differences in infection survival with age among longer-lived mammals; younger seals died of phocine distemper more than subadults [50] and youngest and oldest elephants (but not intermediate ages) were most likely to die from parasite-related causes [51]. Additionally, amphibians, which have distinct life stages, survive infection differently between those stages [52–54]. Here we provide important results showing that younger and older individuals of a wild small mammal have different mortality hazards due to an endemic infection with no obvious pathology, despite their fast life-histories and lack of distinct life stages.

In our study, other host characteristics, and qualities of the host’s environment, affected survival independent of infection. Young, reproductively-active voles had increased survival compared to young non-reproductively active voles. This could be because reproductively active voles maintain territories [42], potentially securing access to resources which are difficult for non-reproductive young voles to obtain. An alternative explanation is that non-reproductive young voles disperse away in search of other habitat (rather than truly dying). Our results suggested that food supplementation decreased survival of voles, however, when transient adult voles were removed from the data set, there was no difference in survival between fed/control and unfed/control voles. This indicates that food may affect movement rather than survival.

Our study addresses a critical need for rigorous field investigations on the impact of pathogens on survival of their reservoir hosts, and key factors potentially modulating that survival. By demonstrating reduced survival of infected reservoir hosts, this study challenges the historical expectation that pathogens with a long coevolutionary relationship do not harm their reservoirs. Importantly, we also show that relationships between pathogens and reservoir host survival can change with host age, with important implications for reservoir infection dynamics.

## Ethics Statement

This research was conducted under approval of the University of Arkansas Institutional Animal Care and Use Committee (IACUC #19105) and the Finnish Animal Ethics Board (ESAVI-17810-2019). Access to field sites was provided by private landowners and by Metsähallitus Metsätalous Oy (MH 6302/2019).

## Conflicts of Interest

The authors report no conflicts of interest.

## Author Contributions

**K.E.W**.: formal analysis, investigation, data curation, visualization, writing - original draft, project administration. **J.S.M.V**.: investigation, data curation, writing - review & editing, project administration. **J.M**.: investigation, data curation, writing - review & editing, project administration. **D.F.H**.: investigation, data curation, writing - review & editing, project administration. **B.B.H**.: conceptualization, formal analysis, writing - review & editing. **C.G.F**.: investigation, writing - review & editing. **T.S**.: methodology, resources, writing - review & editing. **M.E.C**.: conceptualization, methodology, writing - review & editing, funding acquisition. **C.E.C**.: conceptualization, methodology, writing - review & editing, funding acquisition. **R.J.H**.: conceptualization, methodology, writing - review & editing, funding acquisition. **S.A.B**.: conceptualization, methodology, investigation, writing - review & editing, project administration, funding acquisition. **K.M.F**.: conceptualization, methodology, investigation, writing - review & editing, supervision, project administration, funding acquisition.

## Acknowledgements

We are grateful to our collaborators at the Lammi Biological Station and University of Helsinki: John Loehr, Janne Sundell, Esa-Pekka Tuominen, Joni Uusitalo, Tiina Tulonen, Matti Kotakorpi, Riitta Ilola, Jaakko Vainionpää, Tomas Strandin, Mira Utriainen and Leena Palmunen, to the students, research station interns, and collaborators who helped collect and process the field data: Alexis Beagle, Muriel Chaudhri, Stephanie Du, Lucie Fornilli, Mathilde Gaudillère, Teemu Lemola, Nathaniel Mull, Anni Simonen, Shannon Kitchen, Brent Newman, Samuel Clague, Léa Tambareau, Emilie Bonhomme, Eléonore Miston, Amy Schexnayder, Isabella Stark, Stephen Shikaze, Eunice Oh, Juliane Damaschke, Elizabeth Pellegrini, Hannah Chan, Maëlle Pavin De Lafarge, Chéryline Chanrion, Raven Leggett, Anna Bolding, Aniket Shitole, Austin Rife, Oliver Xu, Fiorina Muster, Annabelle Erisman, Mary McFetridge and Amandine Pernot.

## Funding Statement

This research was funded by the National Science Foundation (DEB-1911925) with additional funding to MEC from DEB-2321358.

## Notes

### Competing Interest Statement

The authors have declared no competing interest.

