## Supplemental Figure 1 for "Age-dependent effects of infection on survival of a wild rodent reservoir host"

Fieldwork:

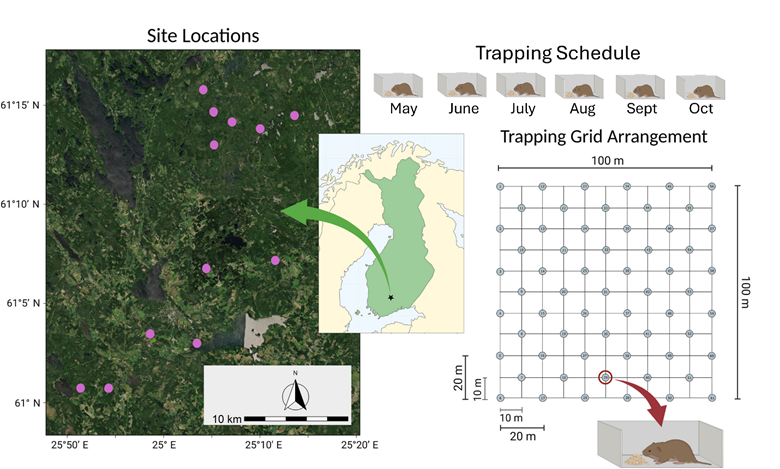

**Supplemental Figure 1:** Longitudinal monitoring methods summary. This figure is adapted from Mistrick et al [1], *J. Anim. Ecol.,* [https://doi.org/10.1111/1365-2656.1406](https://doi.org/10.1111/1365-2656.14067). Licensed under [Creative Commons Attribution](http://creativecommons.org/licenses/by/4.0/).

Deworm treatment and ticks:

To attribute the effects of our deworming medication to nematode infections, we ran a validation analysis exploring potential relationships between deworm treatment and tick infections. One of our deworming medications, Ivermectin, can also treat other parasites found on bank voles, such as ticks [2,3]. During sample collection from all months in 2021, in May through July of 2022, in May and June of 2023, and in the final May trapping in 2024, ticks were counted on the ears and faces of captured voles. This data was used to run a binomial generalized linear mixed model (‘glmmTMB’; [4]) with tick infection as the response variable (binary) and deworm treatment as the explanatory variable (Vole ID was included as a random effect to account for multiple sampling). Similar models were run with tick abundance (the number of ticks counted on a vole’s ears and face, including samples with zero ticks) and intensity of tick infection (the number of ticks counted on a vole’s ears and face, excluding samples with zero ticks) as the response variables.

The binomial tick infection analysis and the tick abundance analysis had a sample size of 1165. There was no significant relationship between deworm treatment and whether or not a vole had ticks (p=0.88), or between deworm treatment and the abundance of ticks on a vole (p=0.28). The abundance model was fit with a Poisson distribution as it was count data. There were 634 samples in the tick infection intensity analysis. The intensity model was fit with a truncated Poisson distribution as it was count data excluding zeros. There was no significant relationship between deworm treatment and tick infection intensity (p=0.14).

Maternal antibodies:

The PUUV infection status (as determined by IFA) of 32 out of ~1500 voles switched from positive to negative during the study. To test if this was due to maternal antibodies, we followed methods by Voutilainen et al [5]. Specifically, we ran a binomial generalized additive mixed model (using R package ‘mgcv’ [6]) with IFA infection status as the response variable, a smoothed mass term as the explanatory variable, and site as a random effect. Probability of IFA infection decreased at first as mass increased, then switched to increasing (Supplemental Figure 2). This shape is what is expected given that very small voles are positive with maternal antibodies, then as they grow, these antibodies wane and true hantavirus infections (and subsequent production of antibodies) begin to increase with increasing mass. The inflection point is the mass below which maternal antibodies may be present. We found the derivative which was closest to zero to determine the inflection point, which was at a mass of 13.8g (Supplemental Figure 2). This analysis suggested that 16 of the 32 voles had maternal antibodies (all 32 voles were removed from the analyses regardless of mass).

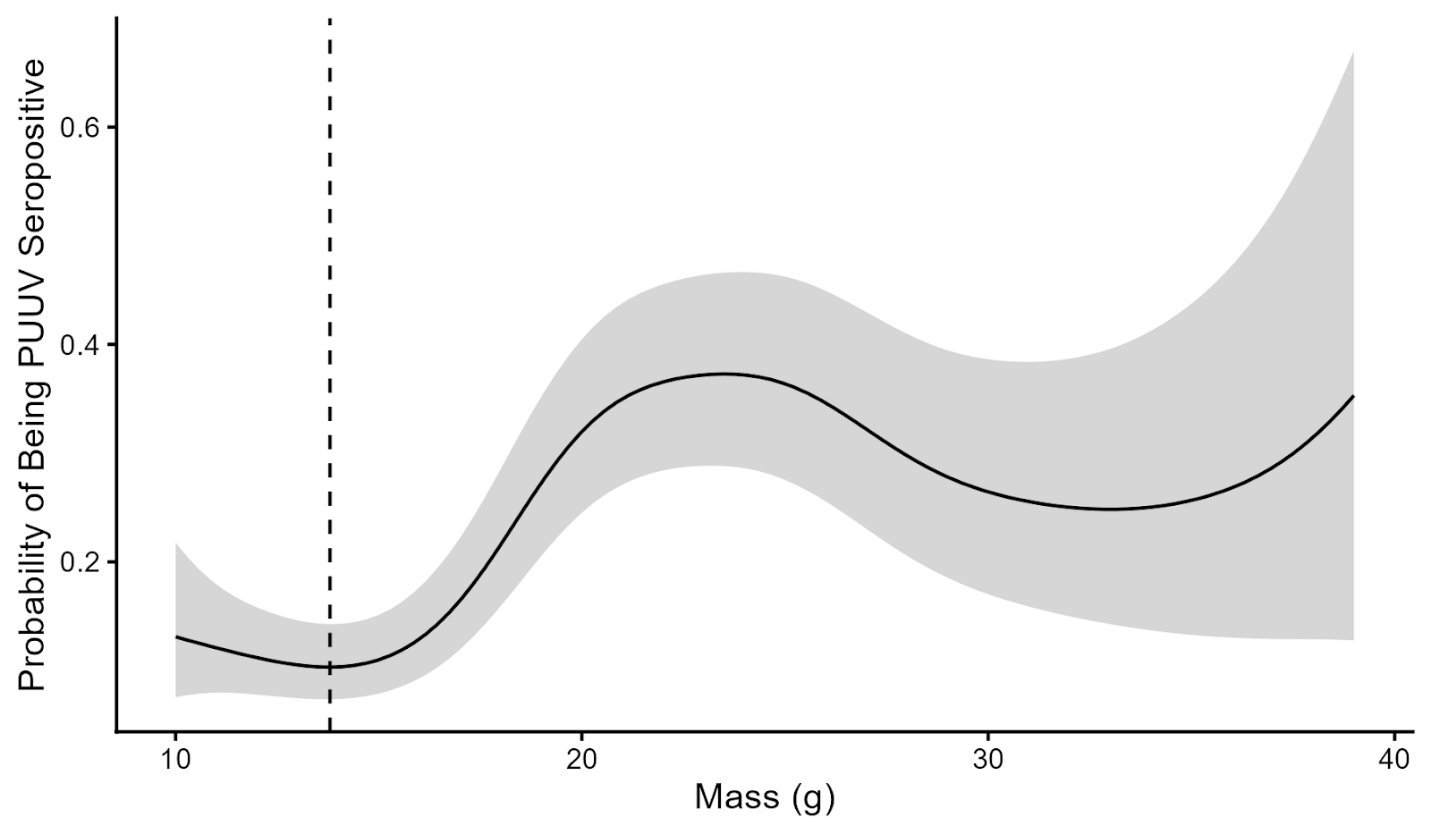

**Supplemental Figure 2:** Predicted results from a GAMM with PUUV antibody status as the response variable and smoothed mass as the explanatory variable. This pattern is what has been shown in previous research and is expected because very small voles have maternal antibodies which wear off as they get larger, and then true infection results in more antibodies among larger voles. The dashed line is where the derivative is closest to 0, indicating the mass at which voles transition from having maternal antibodies to not having maternal antibodies (in this case at 13.8g).

To validate that the results of the primary analysis were not driven by voles with potential maternal antibodies, we removed voles which might have maternal antibodies (i.e., IFA positive and weighing less than 13.8g; Supplemental Figure 2) and reran the primary analysis. However, if any of these small voles had PUUV RNA detected in their saliva, then they were not removed from the analysis as this demonstrates a true infection. This resulted in the removal of 51 samples from the full dataset of 2198 samples (2% removed). The same model was favored in AICc model comparison in the primary analysis with these potential maternal antibody samples removed, and inference from the top model was the same (Supplemental Table 1, Supplemental Table 2). The top model with the highest AICc weight for younger voles in this validation analysis was favored in the main secondary analysis but did not have the highest AICc weight (Supplemental Table 3, Supplemental Table 8). This suggests that the nonsignificant interaction between sex and PUUV infection present in the model with the highest AICc weight in the main secondary analysis may be strengthened by voles with potential maternal antibodies. Importantly, aside from this non-significant interaction, the results for younger voles were the same with and without potential maternal antibodies removed (Supplemental Table 4, Supplemental Table 9). None of the samples from older voles in the secondary analysis were potential maternal antibody samples, so the analysis was identical. These results demonstrate that maternal antibodies were not biasing the results of the primary and secondary analyses.

**Supplemental Table 1**: AICc results from the primary analysis with any measures with potential maternal antibodies removed from the analysis. This model comparison was with the set of Cox proportional hazards models run for the primary analysis. Every model listed included a term clustering by individual vole ID, allowing risk error calculations to account for multiple sampling of the same individuals. The parameters “food” and “deworm” refer to the experimental treatment groups, “puuv” refers to PUUV infection status, “nematode” refers to nematode infection status, and “repro” refers to reproductive status. Any term with “tt()” indicates that this term was allowed to vary with time based on the equation tt=x*time (vole age). These time-varying terms were identified by checking the proportional hazards assumption. The top model with the highest AICc weight (MA1, 0.09 AICc weight) is the same as in the full primary analysis (see Supplemental Table 6).

| **Model Number** | **Model Predictors** | **AICc** | Δ**AICc** | **AICc Wt** |
| --- | --- | --- | --- | --- |
| MA1 | ~puuv + food + deworm + nematode + sex + repro + tt(repro) + month + tt(month) | 8132.43 | 0.00 | 0.09 |
| MA11 | ~nematode*sex + puuv + food + deworm + repro + tt(repro) + month + tt(month) | 8133.04 | 0.61 | 0.07 |
| MA6 | ~deworm*nematode + puuv + food + sex + repro + tt(repro) + month + tt(month) | 8133.22 | 0.79 | 0.06 |
| MA2 | ~puuv*food + deworm + nematode + sex + repro + tt(repro) + month + tt(month) | 8133.29 | 0.86 | 0.06 |
| MA9 | ~food*sex + puuv + deworm + nematode + repro + tt(repro) + month + tt(month) | 8133.73 | 1.30 | 0.05 |
| MA14 | ~puuv*food + nematode*sex + deworm + repro + tt(repro) + month + tt(month) | 8133.83 | 1.40 | 0.05 |
| MA12 | ~puuv*food + deworm*nematode + sex + repro + tt(repro) + month + tt(month) | 8134.09 | 1.66 | 0.04 |
| MA4 | ~puuv*nematode + food + deworm + sex + repro + tt(repro) + month + tt(month) | 8134.40 | 1.98 | 0.03 |
| MA8 | ~nematode*food + puuv + deworm + sex + repro + tt(repro) + month + tt(month) | 8134.42 | 1.99 | 0.03 |
| MA5 | ~puuv*sex + deworm + nematode + food + repro + tt(repro) + month + tt(month) | 8134.44 | 2.01 | 0.03 |
| MA10 | ~deworm*sex + puuv + food + nematode + repro + tt(repro) + month + tt(month) | 8134.44 | 2.02 | 0.03 |
| MA3 | ~puuv*deworm + food + nematode + sex + repro + tt(repro) + month + tt(month) | 8134.44 | 2.02 | 0.03 |
| MA7 | ~food*deworm + puuv + nematode + sex + repro + tt(repro) + month + tt(month) | 8134.45 | 2.02 | 0.03 |
| MA21 | ~deworm*nematode + food*sex + puuv + repro + tt(repro) + month + tt(month) | 8134.54 | 2.12 | 0.03 |
| MA26 | ~nematode*sex + puuv*deworm + food + repro + tt(repro) + month + tt(month) | 8135.05 | 2.62 | 0.03 |
| MA22 | ~food*deworm + nematode*sex + puuv + repro + tt(repro) + month + tt(month) | 8135.05 | 2.62 | 0.03 |
| MA18 | ~puuv*sex + deworm*nematode + food + repro + tt(repro) + month + tt(month) | 8135.24 | 2.81 | 0.02 |
| MA13 | ~puuv*food + deworm*sex + nematode + repro + tt(repro) + month + tt(month) | 8135.31 | 2.88 | 0.02 |
| MA29 | ~puuv*deworm*nematode + food + sex + repro + tt(repro) + month + tt(month) | 8135.66 | 3.23 | 0.02 |
| MA24 | ~food*sex + puuv*nematode + deworm + repro + tt(repro) + month + tt(month) | 8135.72 | 3.29 | 0.02 |
| MA16 | ~puuv*deworm + food*sex + nematode + repro + tt(repro) + month + tt(month) | 8135.75 | 3.32 | 0.02 |
| MA30 | ~puuv*food*sex + deworm + nematode + repro + tt(repro) + month + tt(month) | 8135.89 | 3.46 | 0.02 |
| MA27 | ~puuv*food*deworm + nematode + sex + repro + tt(repro) + month + tt(month) | 8136.09 | 3.66 | 0.01 |
| MA25 | ~deworm*sex + puuv*nematode + food + repro + tt(repro) + month + tt(month) | 8136.42 | 3.99 | 0.01 |
| MA17 | ~puuv*nematode + food*deworm + sex + repro + tt(repro) + month + tt(month) | 8136.42 | 3.99 | 0.01 |
| MA37 | ~puuv*food*deworm + nematode*sex + repro + tt(repro) + month + tt(month) | 8136.44 | 4.01 | 0.01 |
| MA20 | ~puuv*sex + nematode*food + deworm + repro + tt(repro) + month + tt(month) | 8136.44 | 4.01 | 0.01 |
| MA23 | ~nematode*food + deworm*sex + puuv + repro + tt(repro) + month + tt(month) | 8136.44 | 4.01 | 0.01 |
| MA15 | ~puuv*deworm + nematode*food + sex + repro + tt(repro) + month + tt(month) | 8136.44 | 4.01 | 0.01 |
| MA19 | ~puuv*sex + food*deworm + nematode + repro + tt(repro) + month + tt(month) | 8136.46 | 4.03 | 0.01 |
| MA40 | ~puuv*food*sex + deworm*nematode + repro + tt(repro) + month + tt(month) | 8136.59 | 4.16 | 0.01 |
| MA39 | ~puuv*deworm*nematode + food*sex + repro + tt(repro) + month + tt(month) | 8137.08 | 4.65 | 0.01 |
| MA36 | ~food*nematode*sex + puuv + deworm + repro + tt(repro) + month + tt(month) | 8137.10 | 4.67 | 0.01 |
| MA33 | ~food*deworm*nematode + puuv + sex + repro + tt(repro) + month + tt(month) | 8137.26 | 4.83 | 0.01 |
| MA35 | ~deworm*nematode*sex + puuv + food + repro + tt(repro) + month + tt(month) | 8137.91 | 5.48 | 0.01 |
| MA45 | ~deworm*nematode*sex + puuv*food + repro + tt(repro) + month + tt(month) | 8138.67 | 6.25 | 0.00 |
| MA31 | ~puuv*deworm*sex + food + nematode + repro + tt(repro) + month + tt(month) | 8138.69 | 6.26 | 0.00 |
| MA32 | ~puuv*nematode*sex + food + deworm + repro + tt(repro) + month + tt(month) | 8139.03 | 6.60 | 0.00 |
| MA46 | ~food*nematode*sex + puuv*deworm + repro + tt(repro) + month + tt(month) | 8139.12 | 6.69 | 0.00 |
| MA28 | ~puuv*food*nematode + deworm + sex + repro + tt(repro) + month + tt(month) | 8139.18 | 6.75 | 0.00 |
| MA43 | ~food*deworm*nematode + puuv*sex + repro + tt(repro) + month + tt(month) | 8139.28 | 6.85 | 0.00 |
| MA34 | ~food*deworm*sex + puuv + nematode + repro + tt(repro) + month + tt(month) | 8139.65 | 7.23 | 0.00 |
| MA41 | ~puuv*deworm*sex + food*nematode + repro + tt(repro) + month + tt(month) | 8140.68 | 8.25 | 0.00 |
| MA42 | ~puuv*nematode*sex + food*deworm + repro + tt(repro) + month + tt(month) | 8141.05 | 8.62 | 0.00 |
| MA38 | ~puuv*food*nematode + deworm*sex + repro + tt(repro) + month + tt(month) | 8141.19 | 8.77 | 0.00 |
| MA44 | ~food*deworm*sex + puuv*nematode + repro + tt(repro) + month + tt(month) | 8141.65 | 9.22 | 0.00 |
| MA0 | ~1 | 8529.52 | 397.09 | 0.00 |

**Supplemental Table 2:** Inference from MA1, which was favored by AICc in the primary analysis with potential maternal antibody observations removed and had the highest AICc weight (Supplemental Table 1). These results provide information about mortality hazards. The predictors in MA1 were “~puuv + food + deworm + nematode + sex + repro + tt(repro) + month + tt(month)”, and the model was clustered by individual vole ID. In these Cox proportional hazards results, the coefficient gives the direction of the relationship between the explanatory variable and the hazard of dying (positive indicates increased hazard). The exponentiated coefficient gives the hazard ratio and can be used to find the percentage change where 1 is a 0% change in hazard, numbers above 1 are increased hazard (e.g. exp(coef)=1.51 has a 51% higher hazard), and numbers below 1 have decreased hazard (e.g. 0.92 has an 8% decrease in hazard). The relationship with reproductive status changed when voles were 4.3 months old, while the relationship with month changed when voles were 12.2 months old. This timing, and the general pattern of results matches inference from the favored model in the full primary analysis (see Supplemental Table 7), indicating that maternal antibodies are not driving the results.

| **Explanatory variable** | **Coefficient** | **Hazard ratio (exponentiated**  **coefficient)** | **Robust**  **standard error** | **P-value** |
| --- | --- | --- | --- | --- |
| PUUV+ | 0.39 | 1.48 | 0.09 | 2.6e-5*** |
| Fed | 0.16 | 1.17 | 0.07 | 0.029* |
| Deworm | -0.13 | 0.88 | 0.07 | 0.070 |
| Nematode+ | -0.13 | 0.88 | 0.079 | 0.11 |
| Male | 0.22 | 1.25 | 0.07 | 0.0032** |
| Repro active | -1.29 | 0.28 | 0.23 | 2.7e-8*** |
| Time varying repro active | 0.30 | 1.35 | 0.06 | 8.3e-8*** |
| Month | -0.79 | 0.45 | 0.05 | <2e-16*** |
| Time varying month | 0.06 | 1.07 | 0.01 | 3.5e-13*** |

**Supplemental Table 3**: AICc results from the secondary analysis (subsetted into younger and older vole groups) with any measures with potential maternal antibodies removed from the analysis. This model comparison was with the set of Cox proportional hazards models run for the secondary analysis with younger voles (less than 7 months old). Every model listed included a term clustering by individual vole ID, allowing risk error calculations to account for multiple sampling of the same individuals. The parameters “food” and “deworm” refer to the experimental treatment groups, “puuv” refers to PUUV infection status, “nematode” refers to nematode infection status, and “repro” refers to reproductive status. Any term with “tt()” indicates that this term was allowed to vary with time based on the equation tt=x*time (vole age). These time-varying terms were identified by checking the proportional hazards assumption. The top model with the highest AICc weight (MAY1, 0.09 AICc weight) is favored in the main secondary analysis but does not have the highest AICc weight (see Supplemental Table 8). This suggests that the nonsignificant interaction between sex and PUUV infection in MY5 may be strengthened by voles with potential maternal antibodies, but the other results in that model match MAY1 (Supplemental Table 4, Supplemental Table 9).

| **Model Number** | **Model Predictors** | **AICc** | Δ**AICc** | **AICc Wt** |
| --- | --- | --- | --- | --- |
| MAY1 | ~puuv + food + deworm + nematode + sex + repro + tt(repro) + month | 6258.15 | 0.00 | 0.09 |
| MAY5 | ~puuv*sex + deworm + nematode + food + repro + tt(repro) + month | 6258.83 | 0.68 | 0.06 |
| MAY3 | ~puuv*deworm + food + nematode + sex + repro + tt(repro) + month | 6258.96 | 0.81 | 0.06 |
| MAY11 | ~nematode*sex + puuv + food + deworm + repro + tt(repro) + month | 6259.36 | 1.22 | 0.05 |
| MAY4 | ~puuv*nematode + food + deworm + sex + repro + tt(repro) + month | 6259.40 | 1.25 | 0.05 |
| MAY7 | ~food*deworm + puuv + nematode + sex + repro + tt(repro) + month | 6259.66 | 1.51 | 0.04 |
| MAY6 | ~deworm*nematode + puuv + food + sex + repro + tt(repro) + month | 6259.78 | 1.63 | 0.04 |
| MAY2 | ~puuv*food + deworm + nematode + sex + repro + tt(repro) + month | 6259.82 | 1.67 | 0.04 |
| MAY9 | ~food*sex + puuv + deworm + nematode + repro + tt(repro) + month | 6259.93 | 1.79 | 0.04 |
| MAY10 | ~deworm*sex + puuv + food + nematode + repro + tt(repro) + month | 6260.08 | 1.93 | 0.03 |
| MAY8 | ~nematode*food + puuv + deworm + sex + repro + tt(repro) + month | 6260.14 | 1.99 | 0.03 |
| MAY26 | ~nematode*sex + puuv*deworm + food + repro + tt(repro) + month | 6260.17 | 2.02 | 0.03 |
| MAY19 | ~puuv*sex + food*deworm + nematode + repro + tt(repro) + month | 6260.32 | 2.17 | 0.03 |
| MAY18 | ~puuv*sex + deworm*nematode + food + repro + tt(repro) + month | 6260.48 | 2.33 | 0.03 |
| MAY16 | ~puuv*deworm + food*sex + nematode + repro + tt(repro) + month | 6260.72 | 2.58 | 0.03 |
| MAY20 | ~puuv*sex + nematode*food + deworm + repro + tt(repro) + month | 6260.79 | 2.64 | 0.02 |
| MAY17 | ~puuv*nematode + food*deworm + sex + repro + tt(repro) + month | 6260.87 | 2.72 | 0.02 |
| MAY22 | ~food*deworm + nematode*sex + puuv + repro + tt(repro) + month | 6260.88 | 2.73 | 0.02 |
| MAY15 | ~puuv*deworm + nematode*food + sex + repro + tt(repro) + month | 6260.94 | 2.80 | 0.02 |
| MAY24 | ~food*sex + puuv*nematode + deworm + repro + tt(repro) + month | 6261.12 | 2.97 | 0.02 |
| MAY14 | ~puuv*food + nematode*sex + deworm + repro + tt(repro) + month | 6261.12 | 2.97 | 0.02 |
| MAY25 | ~deworm*sex + puuv*nematode + food + repro + tt(repro) + month | 6261.34 | 3.19 | 0.02 |
| MAY12 | ~puuv*food + deworm*nematode + sex + repro + tt(repro) + month | 6261.42 | 3.27 | 0.02 |
| MAY21 | ~deworm*nematode + food*sex + puuv + repro + tt(repro) + month | 6261.57 | 3.42 | 0.02 |
| MAY13 | ~puuv*food + deworm*sex + nematode + repro + tt(repro) + month | 6261.75 | 3.60 | 0.01 |
| MAY29 | ~puuv*deworm*nematode + food + sex + repro + tt(repro) + month | 6261.80 | 3.65 | 0.01 |
| MAY27 | ~puuv*food*deworm + nematode + sex + repro + tt(repro) + month | 6262.03 | 3.88 | 0.01 |
| MAY23 | ~nematode*food + deworm*sex + puuv + repro + tt(repro) + month | 6262.08 | 3.93 | 0.01 |
| MAY33 | ~food*deworm*nematode + puuv + sex + repro + tt(repro) + month | 6262.25 | 4.10 | 0.01 |
| MAY30 | ~puuv*food*sex + deworm + nematode + repro + tt(repro) + month | 6262.38 | 4.23 | 0.01 |
| MAY43 | ~food*deworm*nematode + puuv*sex + repro + tt(repro) + month | 6262.98 | 4.83 | 0.01 |
| MAY31 | ~puuv*deworm*sex + food + nematode + repro + tt(repro) + month | 6263.13 | 4.99 | 0.01 |
| MAY32 | ~puuv*nematode*sex + food + deworm + repro + tt(repro) + month | 6263.20 | 5.05 | 0.01 |
| MAY37 | ~puuv*food*deworm + nematode*sex + repro + tt(repro) + month | 6263.25 | 5.10 | 0.01 |
| MAY39 | ~puuv*deworm*nematode + food*sex + repro + tt(repro) + month | 6263.55 | 5.40 | 0.01 |
| MAY36 | ~food*nematode*sex + puuv + deworm + repro + tt(repro) + month | 6263.61 | 5.46 | 0.01 |
| MAY34 | ~food*deworm*sex + puuv + nematode + repro + tt(repro) + month | 6263.71 | 5.56 | 0.01 |
| MAY40 | ~puuv*food*sex + deworm*nematode + repro + tt(repro) + month | 6264.06 | 5.91 | 0.00 |
| MAY35 | ~deworm*nematode*sex + puuv + food + repro + tt(repro) + month | 6264.41 | 6.26 | 0.00 |
| MAY46 | ~food*nematode*sex + puuv*deworm + repro + tt(repro) + month | 6264.45 | 6.30 | 0.00 |
| MAY28 | ~puuv*food*nematode + deworm + sex + repro + tt(repro) + month | 6264.51 | 6.36 | 0.00 |
| MAY42 | ~puuv*nematode*sex + food*deworm + repro + tt(repro) + month | 6264.65 | 6.50 | 0.00 |
| MAY44 | ~food*deworm*sex + puuv*nematode + repro + tt(repro) + month | 6264.76 | 6.61 | 0.00 |
| MAY41 | ~puuv*deworm*sex + food*nematode + repro + tt(repro) + month | 6265.09 | 6.94 | 0.00 |
| MAY45 | ~deworm*nematode*sex + puuv*food + repro + tt(repro) + month | 6266.10 | 7.95 | 0.00 |
| MAY38 | ~puuv*food*nematode + deworm*sex + repro + tt(repro) + month | 6266.46 | 8.31 | 0.00 |
| MAY0 | ~1 | 6573.91 | 315.76 | 0.00 |

**Supplemental Table 4:** Inference from MAY1, which was favored by AICc  and had the highest AICc weight in the secondary analysis of younger voles with potential maternal antibody observations removed (Supplemental Table 3). The results from this model provide information about mortality hazards. The predictors in MAY1 were “~puuv + food + deworm + nematode + sex + repro + tt(repro) + month”, and clustered by individual vole ID. In these Cox proportional hazards results, the coefficient gives the direction of the relationship between the explanatory variable and the relative hazard of dying (positive indicates increased hazard). The exponentiated coefficient is the hazard ratio and can be used to find the percentage change where 1 is a 0% change in hazard, numbers above 1 are increased hazard (e.g. exp(coef)=1.51 has a 51% higher hazard), and numbers below 1 have decreased hazard (e.g. 0.92 has an 8% decrease in hazard). The relationship with reproductive status changed when voles were 4.3 months old. This timing, and the general pattern of results (besides the lack of a nonsignificant interaction between PUUV and sex) matches inference from the favored model in the full secondary analysis (see Supplemental Table 9), indicating that maternal antibodies are not driving the results.

| **Explanatory variable** | **Coefficient** | **Hazard ratio**  **(exponentiated coefficient)** | **Robust**  **standard error** | **P-value** |
| --- | --- | --- | --- | --- |
| PUUV+ | 0.45 | 1.57 | 0.12 | 0.00015*** |
| Fed | 0.24 | 1.27 | 0.09 | 0.0055** |
| Deworm | -0.23 | 0.80 | 0.08 | 0.0065** |
| Nematode+ | -0.20 | 0.82 | 0.09 | 0.031* |
| Male | 0.20 | 1.22 | 0.09 | 0.020* |
| Repro active | -1.25 | 0.29 | 0.28 | 6.3e-6*** |
| Time varying repro active | 0.29 | 1.34 | 0.08 | 0.00011*** |
| Month | -0.54 | 0.58 | 0.03 | <2e-16*** |

Estimating vole age:

Head-width was used to estimate vole age at first capture to incorporate into the Cox proportional hazards models. Before using the age-head-width relationship described in Kallio et al [7] to determine vole age, we corrected for potential small differences in head-width measurement between researchers. We calculated the mean vole head-width for each month, and the mean head-width for each researcher in that month. The difference between the mean head-width from the researcher and the mean head-width from all researchers was used to adjust each measurement.

General candidate model list:

**Supplemental Table 5:** General list of candidate models used in the analyses. Unless otherwise specified, each analysis used this set of models (with the particular dataset of the analysis) for AICc comparison. For the overwinter analysis, month was not included as it was not a relevant predictor for models specifically examining the likelihood of surviving from October to the following May. Also, in the overwinter analysis, models number 19, 28 and 37 failed to converge, so they were excluded from the AICc comparison to avoid biases arising from different convergences (Supplemental Table 12).

| **Model Number** | **Model Predictors** |
| --- | --- |
| 0 | ~1 |
| 1 | ~puuv + food + deworm + nematode + sex + repro + month |
| 2 | ~puuv*food + deworm + nematode + sex + repro + month |
| 3 | ~puuv*deworm + food + nematode + sex + repro + month |
| 4 | ~puuv*nematode + food + deworm + sex + repro + month |
| 5 | ~puuv*sex + deworm + nematode + food + repro + month |
| 6 | ~deworm*nematode + puuv + food + sex + repro + month |
| 7 | ~food*deworm + puuv + nematode + sex + repro + month |
| 8 | ~nematode*food + puuv + deworm + sex + repro + month |
| 9 | ~food*sex + puuv + deworm + nematode + repro + month |
| 10 | ~deworm*sex + puuv + food + nematode + repro + month |
| 11 | ~nematode*sex + puuv + food + deworm + repro + month |
| 12 | ~puuv*food + deworm*nematode + sex + repro + month |
| 13 | ~puuv*food + deworm*sex + nematode + repro + month |
| 14 | ~puuv*food + nematode*sex + deworm + repro + month |
| 15 | ~puuv*deworm + nematode*food + sex + repro + month |
| 16 | ~puuv*deworm + food*sex + nematode + repro + month |
| 17 | ~puuv*nematode + food*deworm + sex + repro + month |
| 18 | ~puuv*sex + deworm*nematode + food + repro + month |
| 19 | ~puuv*sex + food*deworm + nematode + repro + month |
| 20 | ~puuv*sex + nematode*food + deworm + repro + month |
| 21 | ~deworm*nematode + food*sex + puuv + repro + month |
| 22 | ~food*deworm + nematode*sex + puuv + repro + month |
| 23 | ~nematode*food + deworm*sex + puuv + repro + month |
| 24 | ~food*sex + puuv*nematode + deworm + repro + month |
| 25 | ~deworm*sex + puuv*nematode + food + repro + month |
| 26 | ~nematode*sex + puuv*deworm + food + repro + month |
| 27 | ~puuv*food*deworm + nematode + sex + repro + month |
| 28 | ~puuv*food*nematode + deworm + sex + repro + month |
| 29 | ~puuv*deworm*nematode + food + sex + repro + month |
| 30 | ~puuv*food*sex + deworm + nematode + repro + month |
| 31 | ~puuv*deworm*sex + food + nematode + repro + month |
| 32 | ~puuv*nematode*sex + food + deworm + repro + month |
| 33 | ~food*deworm*nematode + puuv + sex + repro + month |
| 34 | ~food*deworm*sex + puuv + nematode + repro + month |
| 35 | ~deworm*nematode*sex + puuv + food + repro + month |
| 36 | ~food*nematode*sex + puuv + deworm + repro + month |
| 37 | ~puuv*food*deworm + nematode*sex + repro + month |
| 38 | ~puuv*food*nematode + deworm*sex + repro + month |
| 39 | ~puuv*deworm*nematode + food*sex + repro + month |
| 40 | ~puuv*food*sex + deworm*nematode + repro + month |
| 41 | ~puuv*deworm*sex + food*nematode + repro + month |
| 42 | ~puuv*nematode*sex + food*deworm + repro + month |
| 43 | ~food*deworm*nematode + puuv*sex + repro + month |
| 44 | ~food*deworm*sex + puuv*nematode + repro + month |
| 45 | ~deworm*nematode*sex + puuv*food + repro + month |
| 46 | ~food*nematode*sex + puuv*deworm + repro + month |

Vole survival by age:

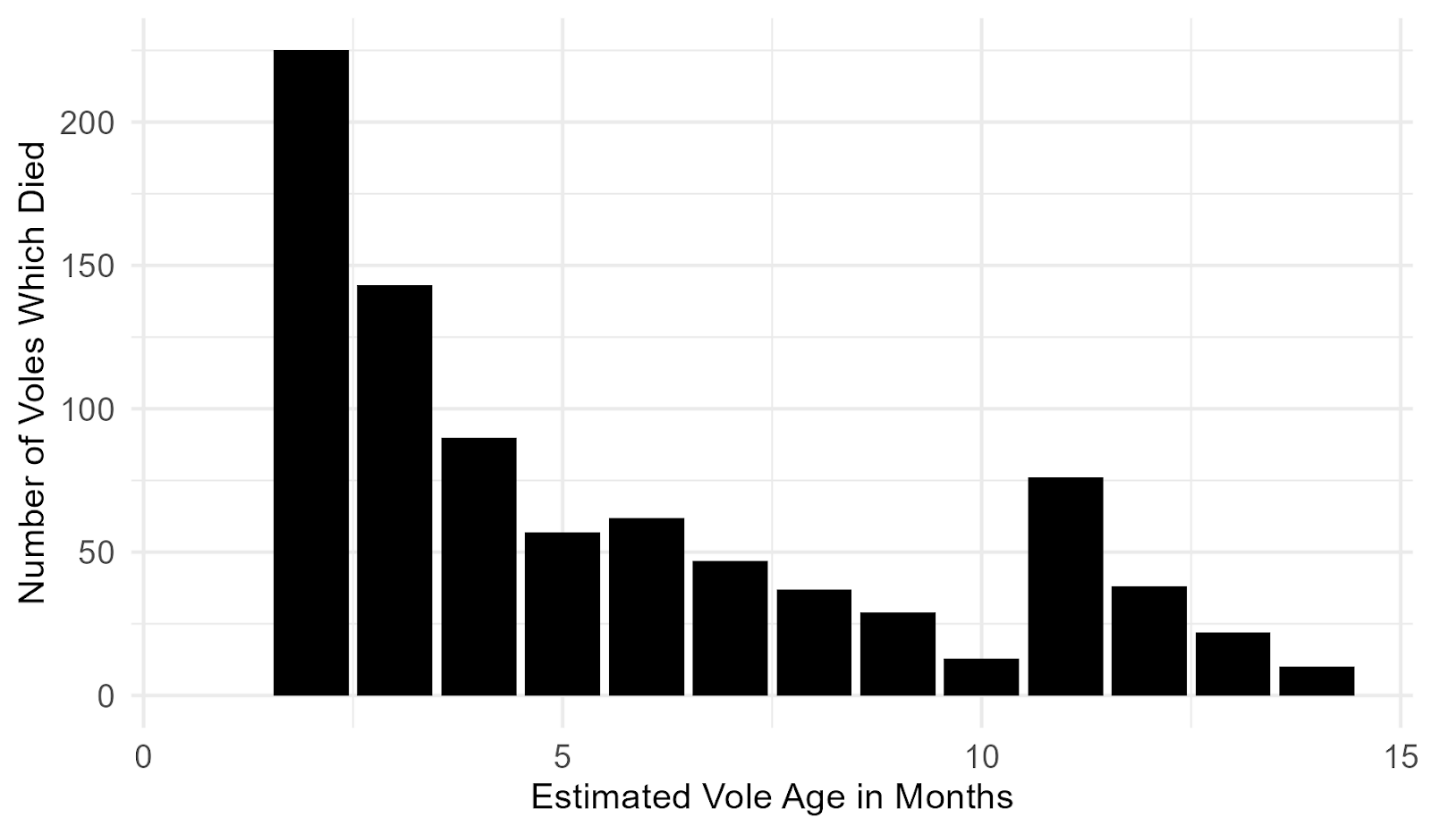

**Supplemental Figure 3:** Visualization of how many voles survived to particular ages (in months) from the raw data. The voles listed at 14 survived for 14 months or longer (the longest surviving vole died at 26 months old). None of the voles that we captured died when they were only one month old (all survived until they were at least 2 months old).

Primary analysis:

**Supplemental Table 6**: AICc results from the primary analysis. This model comparison was with the set of Cox proportional hazards models (Supplemental Table 5). Every model listed included a term clustering by individual vole ID, allowing risk error calculations to account for multiple sampling of the same individuals. The parameters “food” and “deworm” refer to the experimental treatment groups, “puuv” refers to PUUV infection status, “nematode” refers to nematode infection status, and “repro” refers to reproductive status. Any term with “tt()” indicates that this term was allowed to vary with time based on the equation tt=x*time (vole age). These time-varying terms were identified by checking the proportional hazards assumption. The first ten models were favored equally (ΔAICc <2), so the model with the highest AICc weight (M1, 0.10 AICc weight) was used for further inference.

| **Model Number** | **Model Predictors** | **AICc** | Δ**AICc** | **AICc Wt** |
| --- | --- | --- | --- | --- |
| M1 | ~ puuv + food + deworm + nematode + sex + repro + tt(repro) + month + tt(month) | 8584.34 | 0.00 | 0.10 |
| M11 | ~nematode*sex + puuv + food + deworm + repro + tt(repro) + month + tt(month) | 8584.64 | 0.30 | 0.08 |
| M6 | ~deworm*nematode + puuv + food + sex + repro + tt(repro) + month + tt(month) | 8585.37 | 1.03 | 0.06 |
| M9 | ~food*sex + puuv + deworm + nematode + repro + tt(repro) + month + tt(month) | 8585.67 | 1.34 | 0.05 |
| M2 | ~puuv*food + deworm + nematode + sex + repro + tt(repro) + month + tt(month) | 8585.94 | 1.60 | 0.04 |
| M5 | ~puuv*sex + deworm + nematode + food + repro + tt(repro) + month + tt(month) | 8586.00 | 1.67 | 0.04 |
| M3 | ~puuv*deworm + food + nematode + sex + repro + tt(repro) + month + tt(month) | 8586.21 | 1.87 | 0.04 |
| M14 | ~puuv*food + nematode*sex + deworm + repro + tt(repro) + month + tt(month) | 8586.22 | 1.88 | 0.04 |
| M10 | ~deworm*sex + puuv + food + nematode + repro + tt(repro) + month + tt(month) | 8586.27 | 1.94 | 0.04 |
| M7 | ~food*deworm + puuv + nematode + sex + repro + tt(repro) + month + tt(month) | 8586.32 | 1.98 | 0.04 |
| M4 | ~puuv*nematode + food + deworm + sex + repro + tt(repro) + month + tt(month) | 8586.35 | 2.01 | 0.04 |
| M8 | ~nematode*food + puuv + deworm + sex + repro + tt(repro) + month + tt(month) | 8586.35 | 2.02 | 0.04 |
| M26 | ~nematode*sex + puuv*deworm + food + repro + tt(repro) + month + tt(month) | 8586.48 | 2.14 | 0.03 |
| M22 | ~food*deworm + nematode*sex + puuv + repro + tt(repro) + month + tt(month) | 8586.64 | 2.30 | 0.03 |
| M21 | ~deworm*nematode + food*sex + puuv + repro + tt(repro) + month + tt(month) | 8586.73 | 2.39 | 0.03 |
| M12 | ~puuv*food + deworm*nematode + sex + repro + tt(repro) + month + tt(month) | 8586.98 | 2.64 | 0.03 |
| M18 | ~puuv*sex + deworm*nematode + food + repro + tt(repro) + month + tt(month) | 8587.06 | 2.72 | 0.03 |
| M16 | ~puuv*deworm + food*sex + nematode + repro + tt(repro) + month + tt(month) | 8587.57 | 3.23 | 0.02 |
| M24 | ~food*sex + puuv*nematode + deworm + repro + tt(repro) + month + tt(month) | 8587.69 | 3.35 | 0.02 |
| M13 | ~puuv*food + deworm*sex + nematode + repro + tt(repro) + month + tt(month) | 8587.89 | 3.55 | 0.02 |
| M19 | ~puuv*sex + food*deworm + nematode + repro + tt(repro) + month + tt(month) | 8588.02 | 3.65 | 0.02 |
| M20 | ~puuv*sex + nematode*food + deworm + repro + tt(repro) + month + tt(month) | 8588.02 | 3.69 | 0.02 |
| M15 | ~puuv*deworm + nematode*food + sex + repro + tt(repro) + month + tt(month) | 8588.23 | 3.89 | 0.01 |
| M25 | ~deworm*sex + puuv*nematode + food + repro + tt(repro) + month + tt(month) | 8588.29 | 3.95 | 0.01 |
| M23 | ~nematode*food + deworm*sex + puuv + repro + tt(repro) + month + tt(month) | 8588.29 | 3.95 | 0.01 |
| M17 | ~puuv*nematode + food*deworm + sex + repro + tt(repro) + month + tt(month) | 8588.33 | 4.00 | 0.01 |
| M36 | ~food*nematode*sex + puuv + deworm + repro + tt(repro) + month + tt(month) | 8588.45 | 4.11 | 0.01 |
| M30 | ~puuv*food*sex + deworm + nematode + repro + tt(repro) + month + tt(month) | 8588.56 | 4.22 | 0.01 |
| M31 | ~puuv*deworm*sex + food + nematode + repro + tt(repro) + month + tt(month) | 8588.56 | 4.23 | 0.01 |
| M29 | ~puuv*deworm*nematode + food + sex + repro + tt(repro) + month + tt(month) | 8589.09 | 4.75 | 0.01 |
| M33 | ~food*deworm*nematode + puuv + sex + repro + tt(repro) + month + tt(month) | 8589.51 | 5.18 | 0.01 |
| M40 | ~puuv*food*sex + deworm*nematode + repro + tt(repro) + month + tt(month) | 8589.54 | 5.20 | 0.01 |
| M27 | ~puuv*food*deworm + nematode + sex + repro + tt(repro) + month + tt(month) | 8589.61 | 5.28 | 0.01 |
| M37 | ~puuv*food*deworm + nematode*sex + repro + tt(repro) + month + tt(month) | 8589.68 | 5.34 | 0.01 |
| M35 | ~deworm*nematode*sex + puuv + food + repro + tt(repro) + month + tt(month) | 8589.73 | 5.39 | 0.01 |
| M32 | ~puuv*nematode*sex + food + deworm + repro + tt(repro) + month + tt(month) | 8590.22 | 5.88 | 0.01 |
| M46 | ~food*nematode*sex + puuv*deworm + repro + tt(repro) + month + tt(month) | 8590.27 | 5.93 | 0.01 |
| M39 | ~puuv*deworm*nematode + food*sex + repro + tt(repro) + month + tt(month) | 8590.56 | 6.22 | 0.00 |
| M41 | ~puuv*deworm*sex + food*nematode + repro + tt(repro) + month + tt(month) | 8590.58 | 6.25 | 0.00 |
| M43 | ~food*deworm*nematode + puuv*sex + repro + tt(repro) + month + tt(month) | 8591.27 | 6.93 | 0.00 |
| M45 | ~deworm*nematode*sex + puuv*food + repro + tt(repro) + month + tt(month) | 8591.30 | 6.97 | 0.00 |
| M34 | ~food*deworm*sex + puuv + nematode + repro + tt(repro) + month + tt(month) | 8591.41 | 7.07 | 0.00 |
| M28 | ~puuv*food*nematode + deworm + sex + repro + tt(repro) + month + tt(month) | 8591.58 | 7.24 | 0.00 |
| M42 | ~puuv*nematode*sex + food*deworm + repro + tt(repro) + month + tt(month) | 8592.21 | 7.88 | 0.00 |
| M44 | ~food*deworm*sex + puuv*nematode + repro + tt(repro) + month + tt(month) | 8593.43 | 9.09 | 0.00 |
| M38 | ~puuv*food*nematode + deworm*sex + repro + tt(repro) + month + tt(month) | 8593.55 | 9.21 | 0.00 |
| M0 | ~1 | 9006.60 | 422.26 | 0.00 |

**Supplemental Table 7:** Inference from M1, which was favored by AICc and had the highest AICc weight (Supplemental Table 6). The results from this model provide information about mortality hazards. The predictors in M1 were “~puuv + food + deworm + nematode + sex + repro + tt(repro) + month + tt(month)”, and the model was clustered by individual vole ID. In these Cox proportional hazards results, the coefficient gives the direction of the relationship between the explanatory variable and the relative hazard of dying (positive indicates increased hazard). The exponentiated coefficient is the hazard ratio and can be used to find the percentage change where 1 is a 0% change in hazard, numbers above 1 are increased hazard (e.g. exp(coef)=1.51 has a 51% higher hazard), and numbers below 1 have decreased hazard (e.g. 0.92 has an 8% decrease in hazard). The relationship with reproductive status changed when voles were 4.3 months old, while the relationship with month changed when voles were 12.4 months old.

| **Explanatory variable** | **Coefficient** | **Hazard ratio (exponentiated coefficient)** | **Robust standard error** | **P-value** |
| --- | --- | --- | --- | --- |
| PUUV + | 0.47 | 1.59 | 0.08 | 2.3e-8*** |
| Fed | 0.17 | 1.18 | 0.07 | 0.018* |
| Deworm | -0.12 | 0.89 | 0.07 | 0.085 |
| Nematode+ | -0.13 | 0.88 | 0.08 | 0.10 |
| Male | 0.20 | 1.22 | 0.07 | 0.0060** |
| Repro active | -1.26 | 0.28 | 0.23 | 2.2e-8*** |
| Time varying repro active | 0.29 | 1.34 | 0.05 | 1.13e-7*** |
| Month | -0.77 | 0.46 | 0.05 | <2e-16*** |
| Time varying month | 0.06 | 1.06 | 0.01 | 1.2e-12*** |

Secondary age analysis:

**Supplemental Table 8**: AICc results from the secondary analysis, dividing the analysis into vole age groups. This model comparison was with the set of Cox proportional hazards models (Supplemental Table 5) run on data from voles less than 7 months old (younger voles). Every model listed included a term clustering by individual vole ID, allowing risk error calculations to account for multiple sampling of the same individuals. The parameters “food” and “deworm” refer to the experimental treatment groups, “puuv” refers to PUUV infection status, “nematode” refers to nematode infection status, and “repro” refers to reproductive status. Any term with “tt()” indicates that this term was allowed to vary with time based on the equation tt=x*time (vole age). These time-varying terms were identified by checking the proportional hazards assumption. The first nine models were favored equally (ΔAICc <2). MY5 had the highest AICc weight (0.10 AICc weight) and was used for further inference.

| **Model Number** | **Model Predictors** | **AICc** | Δ**AICc** | **AICc Wt** |
| --- | --- | --- | --- | --- |
| MY5 | ~puuv*sex + deworm + nematode + food + repro + tt(repro) + month | 6707.27 | 0.00 | 0.10 |
| MY1 | ~puuv + food + deworm + nematode + sex + repro + tt(repro) + month | 6707.65 | 0.37 | 0.08 |
| MY19 | ~puuv*sex + food*deworm + nematode + repro + tt(repro) + month | 6708.47 | 1.20 | 0.05 |
| MY11 | ~nematode*sex + puuv + food + deworm + repro + tt(repro) + month | 6708.55 | 1.28 | 0.05 |
| MY18 | ~puuv*sex + deworm*nematode + food + repro + tt(repro) +month | 6708.87 | 1.59 | 0.04 |
| MY7 | ~food*deworm + puuv + nematode + sex + repro + tt(repro) + month | 6708.90 | 1.62 | 0.04 |
| MY4 | ~puuv*nematode + food + deworm + sex + repro + tt(repro) + month | 6709.11 | 1.84 | 0.04 |
| MY6 | ~deworm*nematode + puuv + food + sex + repro + tt(repro) + month | 6709.19 | 1.91 | 0.04 |
| MY2 | ~puuv*food + deworm + nematode + sex + repro + tt(repro) + month | 6709.21 | 1.94 | 0.04 |
| MY20 | ~puuv*sex + nematode*food + deworm + repro + tt(repro) + month | 6709.29 | 2.02 | 0.04 |
| MY10 | ~deworm*sex + puuv + food + nematode + repro + tt(repro) + month | 6709.33 | 2.06 | 0.04 |
| MY9 | ~food*sex + puuv + deworm + nematode + repro + tt(repro) + month | 6709.45 | 2.18 | 0.03 |
| MY3 | ~puuv*deworm + food + nematode + sex + repro + tt(repro) + month | 6709.51 | 2.24 | 0.03 |
| MY8 | ~nematode*food + puuv + deworm + sex + repro + tt(repro) + month | 6709.66 | 2.39 | 0.03 |
| MY22 | ~food*deworm + nematode*sex + puuv + repro + tt(repro) + month | 6709.82 | 2.55 | 0.03 |
| MY14 | ~puuv*food + nematode*sex + deworm + repro + tt(repro) + month | 6710.10 | 2.92 | 0.02 |
| MY17 | ~puuv*nematode + food*deworm + sex + repro + tt(repro) + month | 6710.34 | 3.07 | 0.02 |
| MY26 | ~nematode*sex + puuv*deworm + food + repro + tt(repro) + month | 6710.40 | 3.12 | 0.02 |
| MY12 | ~puuv*food + deworm*nematode + sex + repro + tt(repro) + month | 6710.71 | 3.44 | 0.02 |
| MY25 | ~deworm*sex + puuv*nematode + food + repro + tt(repro) + month | 6710.82 | 3.55 | 0.02 |
| MY13 | ~puuv*food + deworm*sex + nematode + repro + tt(repro) + month | 6710.88 | 3.61 | 0.02 |
| MY24 | ~food*sex + puuv*nematode + deworm + repro + tt(repro) + month | 6710.89 | 3.62 | 0.02 |
| MY21 | ~deworm*nematode + food*sex + puuv + repro + tt(repro) + month | 6711.00 | 3.72 | 0.02 |
| MY31 | ~puuv*deworm*sex + food + nematode + repro + tt(repro) + month | 6711.30 | 4.02 | 0.01 |
| MY16 | ~puuv*deworm + food*sex + nematode + repro + tt(repro) + month | 6711.31 | 4.04 | 0.01 |
| MY32 | ~puuv*nematode*sex + food + deworm + repro + tt(repro) + month | 6711.35 | 4.07 | 0.01 |
| MY23 | ~nematode*food + deworm*sex + puuv + repro + tt(repro) + month | 6711.35 | 4.08 | 0.01 |
| MY30 | ~puuv*food*sex + deworm + nematode + repro + tt(repro) + month | 6711.47 | 4.20 | 0.01 |
| MY15 | ~puuv*deworm + nematode*food + sex + repro + tt(repro) + month | 6711.53 | 4.26 | 0.01 |
| MY43 | ~food*deworm*nematode + puuv*sex + repro + tt(repro) + month | 6711.65 | 4.38 | 0.01 |
| MY33 | ~food*deworm*nematode + puuv + sex + repro + tt(repro) + month | 6711.85 | 4.57 | 0.01 |
| MY36 | ~food*nematode*sex + puuv + deworm + repro + tt(repro) + month | 6712.47 | 5.20 | 0.01 |
| MY34 | ~food*deworm*sex + puuv + nematode + repro + tt(repro) + month | 6712.52 | 5.25 | 0.01 |
| MY42 | ~puuv*nematode*sex + food*deworm + repro + tt(repro) + month | 6712.57 | 5.30 | 0.01 |
| MY40 | ~puuv*food*sex + deworm*nematode + repro + tt(repro) + month | 6713.10 | 5.82 | 0.01 |
| MY35 | ~deworm*nematode*sex + puuv + food + repro + tt(repro) + month | 6713.15 | 5.88 | 0.01 |
| MY41 | ~puuv*deworm*sex + food*nematode + repro + tt(repro) + month | 6713.32 | 6.04 | 0.00 |
| MY27 | ~puuv*food*deworm + nematode + sex + repro + tt(repro) + month | 6713.51 | 6.24 | 0.00 |
| MY29 | ~puuv*deworm*nematode + food + sex + repro + tt(repro) + month | 6713.66 | 6.39 | 0.00 |
| MY44 | ~food*deworm*sex + puuv*nematode + repro + tt(repro) + month | 6713.85 | 6.58 | 0.00 |
| MY28 | ~puuv*food*nematode + deworm + sex + repro + tt(repro) + month | 6713.89 | 6.61 | 0.00 |
| MY46 | ~food*nematode*sex + puuv*deworm + repro + tt(repro) + month | 6714.36 | 7.08 | 0.00 |
| MY37 | ~puuv*food*deworm + nematode*sex + repro + tt(repro) + month | 6714.40 | 7.13 | 0.00 |
| MY45 | ~deworm*nematode*sex + puuv*food + repro + tt(repro) + month | 6714.69 | 7.42 | 0.00 |
| MY39 | ~puuv*deworm*nematode + food*sex + repro + tt(repro) + month | 6715.48 | 8.21 | 0.00 |
| MY38 | ~puuv*food*nematode + deworm*sex + repro + tt(repro) + month | 6715.62 | 8.35 | 0.00 |
| MY0 | ~1 | 7050.98 | 343.71 | 0.00 |

**Supplemental Table 9:** Inference for the younger and older vole analyses. Candidate models run in the primary analysis were run for younger voles (less than 7 months) or older voles (7 months or older). The results from these models provide information about mortality hazards. For younger voles, MY5 was favored with the highest AICc weight. The predictors in MY5 were “~puuv*sex + deworm + nematode + food + repro + tt(repro) + month” and the model was clustered by vole ID. The results of MY5 are shown here. For older voles, both MO21 (model predictors: “~deworm*nematode + food*sex + puuv + repro + month”) and MO9 (model predictors: “~food*sex + puuv + deworm + nematode + repro + month”) were favored with the highest AICc weight. The results from MO9 are shown here, while the results of MO21 can be found in Supplemental Table 11. In these results, the coefficient gives the direction of the relationship between the explanatory variable and the relative hazard of dying (positive indicates increased hazard). The exponentiated coefficient is the hazard ratio and can be used to find the percentage change where 1 is a 0% change in hazard, numbers above 1 are increased hazard (e.g. exp(coef)=1.51 has a 51% higher hazard), and numbers below 1 have decreased hazard (e.g. 0.92 has an 8% decrease in hazard). In MY5, the relationship with reproductive status changed direction when voles were 4.4 months old. See Supplemental Figure 4 for interpretation of the Male*PUUV+ interaction in MY5 and Supplemental Figure 5 for interpretation of the Male*Fed interaction in MO9.

**Younger voles:**

| **Explanatory variable** | **Coefficient** | **Hazard ratio (exponentiated**  **coefficient)** | **Robust**  **standard error** | **P-value** |
| --- | --- | --- | --- | --- |
| PUUV + | 0.75 | 2.11 | 0.15 | 1.0e-6*** |
| Fed | 0.25 | 1.28 | 0.08 | 0.0036** |
| Deworm | -0.20 | 0.81 | 0.08 | 0.011* |
| Nematode+ | -0.20 | 0.82 | 0.09 | 0.029* |
| Male | 0.24 | 1.27 | 0.09 | 0.0099** |
| Repro active | -1.22 | 0.30 | 0.27 | 6.6e-6*** |
| Time varying repro active | 0.28 | 1.32 | 0.07 | 0.00016*** |
| Month | -0.54 | 0.58 | 0.03 | <2e-16*** |
| Male*PUUV+ | -0.33 | 0.72 | 0.19 | 0.093 |

**Older voles:**

| **Explanatory variable** | **Coefficient** | **Hazard ratio**  **(exponentiated coefficient)** | **Robust**  **standard error** | **P-value** |
| --- | --- | --- | --- | --- |
| PUUV + | 0.13 | 1.14 | 0.14 | 0.36 |
| Fed | -0.43 | 0.65 | 0.23 | 0.06 |
| Deworm | 0.09 | 1.10 | 0.13 | 0.48 |
| Nematode+ | -0.07 | 0.93 | 0.14 | 0.59 |
| Male | 0.22 | 1.24 | 0.19 | 0.25 |
| Repro active | 1.37 | 3.94 | 0.39 | 0.00037*** |
| Month | 0.06 | 1.06 | 0.07 | 0.36 |
| Male*Fed | 0.44 | 1.55 | 0.28 | 0.11 |

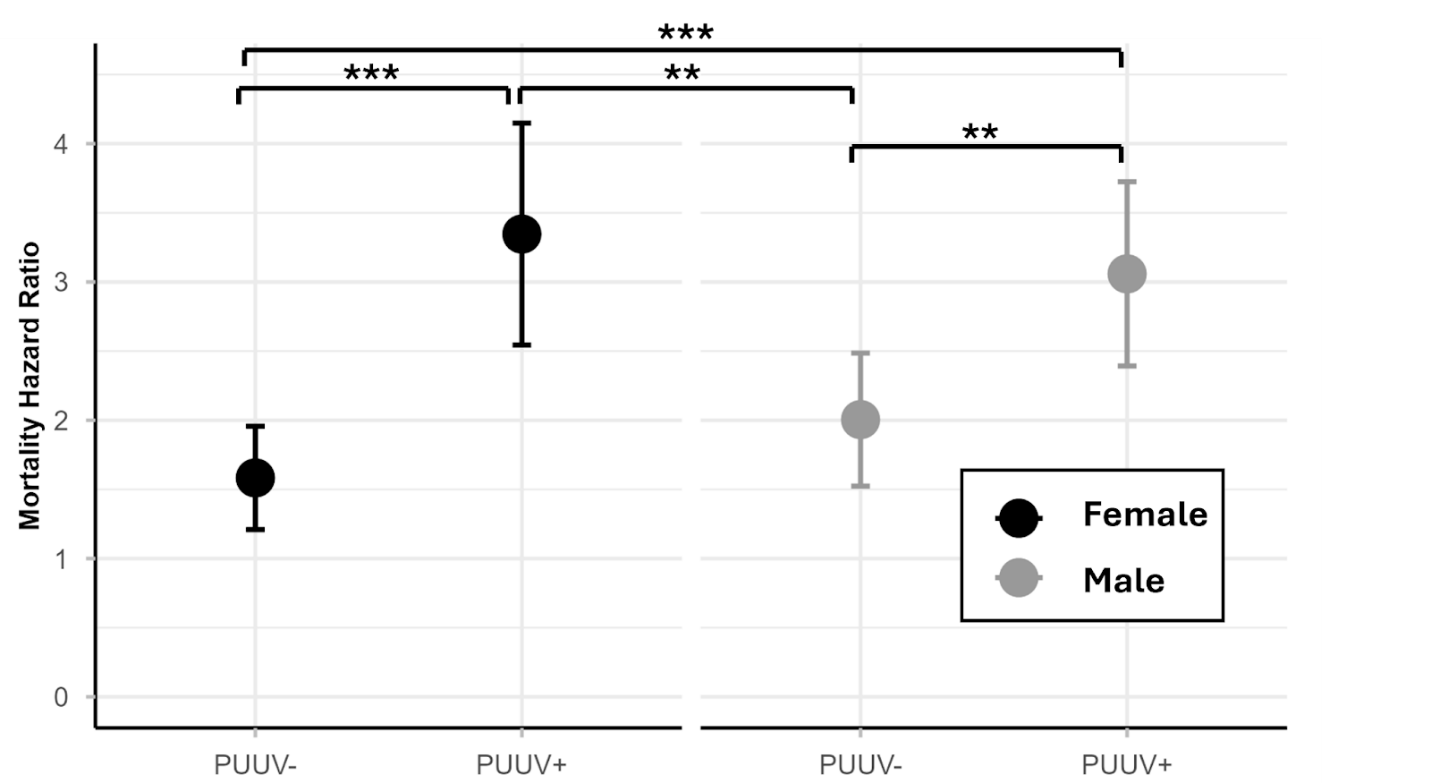

**Supplemental Figure 4**: Plot of nonsignificant interaction between PUUV and sex in MY5 (the top favored model with the highest AICc weight). This plot shows predicted mortality hazard ratios, relative to the sample average for all predictors in the Cox proportional hazards model (MY5). This illustrates that in every comparison, PUUV infected voles had a higher relative mortality hazard than PUUV uninfected voles. Significance was calculated from post hoc pairwise comparisons (with Bonferroni correction).

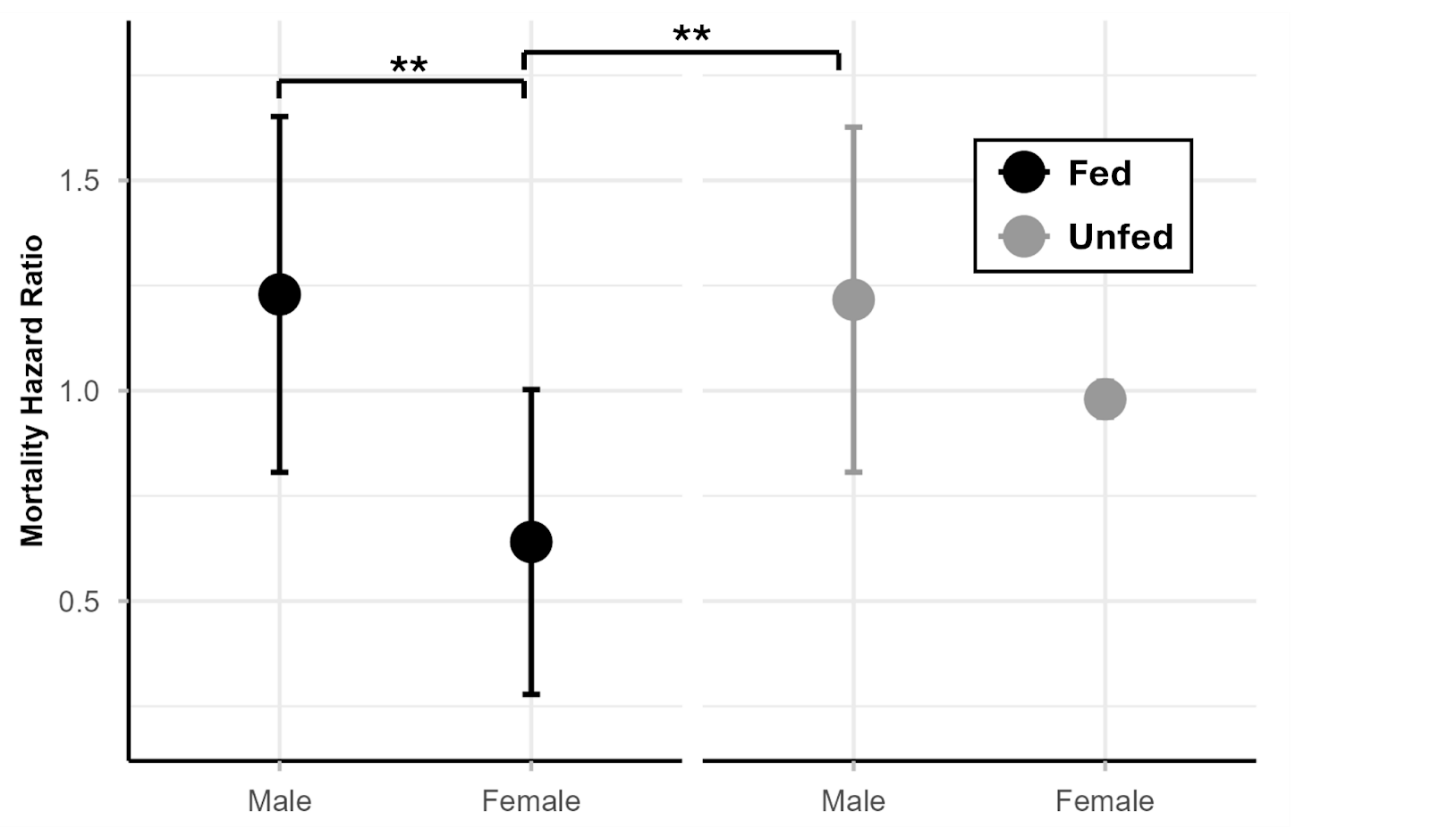

**Supplemental Figure 5:** Plot of nonsignificant interaction between food treatment and sex in MO9 (one of the top favored models with the highest AICc weight in the older vole secondary analysis). This plot shows predicted mortality hazard ratios, relative to the sample average for all predictors in the Cox proportional hazards model (MO9). We detected that males have higher relative mortality hazard than females when receiving food treatment, but not at control sites. Significance was calculated from post hoc pairwise comparisons (with Bonferroni correction).

**Supplemental Table 10**: AICc results from the secondary analysis, dividing the analysis into vole age groups. This model comparison was with the set of Cox proportional hazards models (Supplemental Table 5) run on data from voles 7 months old or older (older voles). Every model listed included a term clustering by individual vole ID, allowing risk error calculations to account for multiple sampling of the same individuals. The parameters “food” and “deworm” refer to the experimental treatment groups, “puuv” refers to PUUV infection status, “nematode” refers to nematode infection status, and “repro” refers to reproductive status. There were no time-varying terms identified in this group of models. The first 14 models were favored equally (ΔAICc <2). MO9 and MO21 had the highest AICc weight (0.07 AICc weight) so they were used for further inference.

| **Model Number** | **Model Predictors** | **AICc** | Δ**AICc** | **AICc Wt** |
| --- | --- | --- | --- | --- |
| MO9 | ~food*sex + puuv + deworm + nematode + repro + month | 1835.49 | 0.00 | 0.07 |
| MO21 | ~deworm*nematode + food*sex + puuv + repro + month | 1835.58 | 0.10 | 0.07 |
| MO6 | ~deworm*nematode + puuv + food + sex + repro + month | 1835.64 | 0.15 | 0.06 |
| MO1 | ~puuv + food + deworm + nematode + sex + repro + month | 1835.66 | 0.17 | 0.06 |
| MO24 | ~food*sex + puuv*nematode + deworm + repro + month | 1835.70 | 0.22 | 0.06 |
| MO4 | ~puuv*nematode + food + deworm + sex + repro + month | 1836.03 | 0.54 | 0.05 |
| MO7 | ~food*deworm + puuv + nematode + sex + repro + month | 1836.42 | 0.94 | 0.04 |
| MO11 | ~nematode*sex + puuv + food + deworm + repro + month | 1836.49 | 1.00 | 0.04 |
| MO2 | ~puuv*food + deworm + nematode + sex + repro + month | 1836.93 | 1.45 | 0.03 |
| MO17 | ~puuv*nematode + food*deworm + sex + repro + month | 1836.97 | 1.49 | 0.03 |
| MO12 | ~puuv*food + deworm*nematode + sex + repro + month | 1837.05 | 1.56 | 0.03 |
| MO22 | ~food*deworm + nematode*sex + puuv + repro + month | 1837.18 | 1.70 | 0.03 |
| MO8 | ~nematode*food + puuv + deworm + sex + repro + month | 1837.46 | 1.97 | 0.03 |
| MO10 | ~deworm*sex + puuv + food + nematode + repro + month | 1837.48 | 1.99 | 0.03 |
| MO16 | ~puuv*deworm + food*sex + nematode + repro + month | 1837.49 | 2.01 | 0.03 |
| MO18 | ~puuv*sex + deworm*nematode + food + repro + month | 1837.51 | 2.03 | 0.03 |
| MO5 | ~puuv*sex + deworm + nematode + food + repro + month | 1837.63 | 2.14 | 0.02 |
| MO25 | ~deworm*sex + puuv*nematode + food + repro + month | 1837.64 | 2.16 | 0.02 |
| MO3 | ~puuv*deworm + food + nematode + sex + repro + month | 1837.65 | 2.16 | 0.02 |
| MO14 | ~puuv*food + nematode*sex + deworm + repro + month | 1837.87 | 2.38 | 0.02 |
| MO19 | ~puuv*sex + food*deworm + nematode + repro + month | 1838.41 | 2.93 | 0.02 |
| MO30 | ~puuv*food*sex + deworm + nematode + repro + month | 1838.49 | 3.01 | 0.02 |
| MO26 | ~nematode*sex + puuv*deworm + food + repro + month | 1838.54 | 3.05 | 0.02 |
| MO40 | ~puuv*food*sex + deworm*nematode + repro + month | 1838.63 | 3.14 | 0.01 |
| MO34 | ~food*deworm*sex + puuv + nematode + repro + month | 1838.68 | 3.19 | 0.01 |
| MO13 | ~puuv*food + deworm*sex + nematode + repro + month | 1838.78 | 3.29 | 0.01 |
| MO44 | ~food*deworm*sex + puuv*nematode + repro + month | 1838.87 | 3.38 | 0.01 |
| MO31 | ~puuv*deworm*sex + food + nematode + repro + month | 1839.26 | 3.77 | 0.01 |
| MO23 | ~nematode*food + deworm*sex + puuv + repro + month | 1839.30 | 3.82 | 0.01 |
| MO20 | ~puuv*sex + nematode*food + deworm + repro + month | 1839.43 | 3.95 | 0.01 |
| MO36 | ~food*nematode*sex + puuv + deworm + repro + month | 1839.45 | 3.97 | 0.01 |
| MO15 | ~puuv*deworm + nematode*food + sex + repro + month | 1839.47 | 3.98 | 0.01 |
| MO39 | ~puuv*deworm*nematode + food*sex + repro + month | 1839.74 | 4.25 | 0.01 |
| MO29 | ~puuv*deworm*nematode + food + sex + repro + month | 1839.92 | 4.43 | 0.01 |
| MO35 | ~deworm*nematode*sex + puuv + food + repro + month | 1840.15 | 4.67 | 0.01 |
| MO33 | ~food*deworm*nematode + puuv + sex + repro + month | 1840.47 | 4.99 | 0.01 |
| MO41 | ~puuv*deworm*sex + food*nematode + repro + month | 1841.07 | 5.58 | 0.00 |
| MO28 | ~puuv*food*nematode + deworm + sex + repro + month | 1841.47 | 5.99 | 0.00 |
| MO32 | ~puuv*nematode*sex + food + deworm + repro + month | 1841.51 | 6.03 | 0.00 |
| MO46 | ~food*nematode*sex + puuv*deworm + repro + month | 1841.56 | 6.07 | 0.00 |
| MO45 | ~deworm*nematode*sex + puuv*food + repro + month | 1841.65 | 6.16 | 0.00 |
| MO27 | ~puuv*food*deworm + nematode + sex + repro + month | 1841.99 | 6.51 | 0.00 |
| MO42 | ~puuv*nematode*sex + food*deworm + repro + month | 1842.40 | 6.91 | 0.00 |
| MO43 | ~food*deworm*nematode + puuv*sex + repro + month | 1842.42 | 6.93 | 0.00 |
| MO37 | ~puuv*food*deworm + nematode*sex + repro + month | 1842.87 | 7.38 | 0.00 |
| MO38 | ~puuv*food*nematode + deworm*sex + repro + month | 1843.20 | 7.71 | 0.00 |
| MO0 | ~1 | 1855.82 | 20.34 | 0.00 |

**Supplemental Table 11:** Inference from MO21, one model favored with the highest AICc weight in the older vole models from the secondary analysis. The results from this model provide information about mortality hazards. The predictors in MO21 were “~deworm*nematode + food*sex + puuv + repro + month” and the model was clustered by vole ID. In these results, the coefficient gives the direction of the relationship between the explanatory variable and the relative hazard of dying (positive indicates increased hazard). The exponentiated coefficient is the hazard ratio and can be used to find the percentage change where 1 is a 0% change in hazard, numbers above 1 are increased hazard (e.g. exp(coef)=1.51 has a 51% higher hazard), and numbers below 1 have decreased hazard (e.g. 0.92 has an 8% decrease in hazard). See Supplemental Figure 6 for interpretation of the Male*Fed interaction in MO21. A post hoc comparison (Bonferroni corrected) confirmed that there were no significant differences between any pairs of predictors in the Nematode+*Deworm interaction in MO21.

| **Explanatory variable** | **Coefficient** | **Hazard ratio (exponentiated**  **coefficient)** | **Robust**  **standard error** | **P-value** |
| --- | --- | --- | --- | --- |
| PUUV + | 0.14 | 1.15 | 0.14 | 0.31 |
| Fed | -0.43 | 0.65 | 0.23 | 0.059 |
| Deworm | -0.08 | 0.92 | 0.18 | 0.64 |
| Nematode+ | -0.26 | 0.77 | 0.19 | 0.18 |
| Male | 0.22 | 1.25 | 0.19 | 0.24 |
| Repro active | 1.36 | 3.88 | 0.38 | 0.00037*** |
| Month | 0.06 | 1.06 | 0.07 | 0.36 |
| Male*Fed | 0.43 | 1.53 | 0.27 | 0.12 |
| Deworm*Nematode+ | 0.38 | 1.47 | 0.27 | 0.16 |

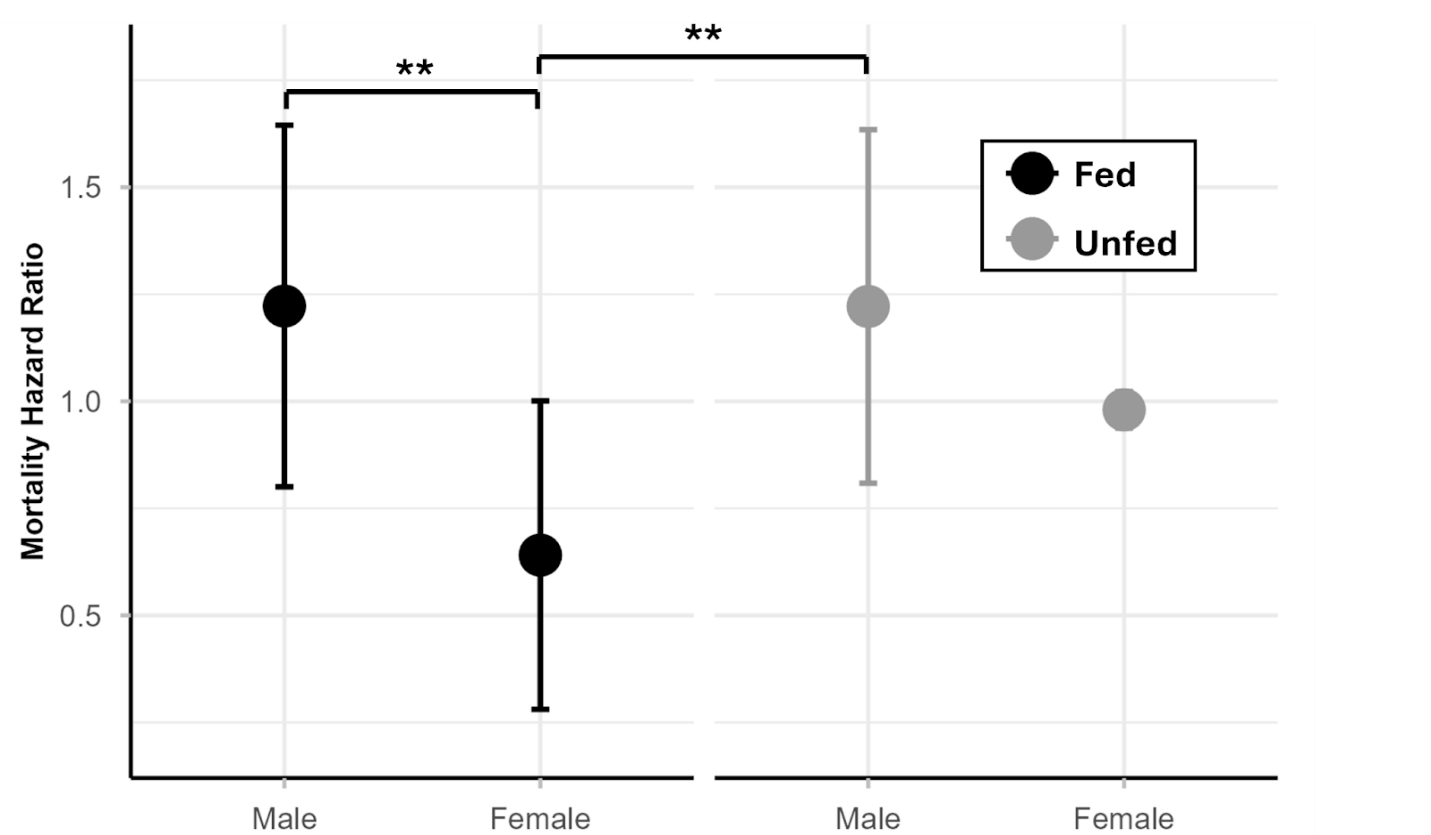

**Supplemental Figure 6:** Plot of nonsignificant interaction between food treatment and sex in MO21 (one of the top favored models with the highest AICc weight in the older vole secondary analysis). This plot shows predicted mortality hazard ratios, relative to the sample average for all predictors in the Cox proportional hazards model (MO21). We detected that males have higher relative mortality hazard than females when receiving food treatment, but not at control sites. Significance was calculated from post hoc pairwise comparisons (with Bonferroni correction).

Overwinter survival analysis:

**Supplemental Table 12:** AICc results from the overwinter survival analysis. These were GLMM models with binomial error distribution (since the response variable was a binary variable of whether or not the vole survived the winter), and every model included a random effect of site to account for inherent site differences. These were the same candidate models listed in Supplemental Table 5, but without a “month” covariate, as that is not relevant to the overwinter survival analysis. Additionally, 3 models (W19, W28, and W37) did not converge, and were excluded from the model comparison to avoid biasing the AICc with differing convergences. The parameters “food” and “deworm” refer to the experimental treatment groups, “puuv” refers to PUUV infection status in October, “nematode” refers to nematode infection status in October, and “repro” refers to reproductive status. No model was favored over the null model (all of the favored models were within 2 units of the null model), indicating that none of the predictors helped to explain the variance in overwinter survival. As such, no further inference was made on the overwinter survival models.

| **Model Number** | **Model Predictors** | **AICc** | Δ**AICc** | **AICc Wt** |
| --- | --- | --- | --- | --- |
| W5 | ~puuv*sex + deworm + nematode + food + repro | 232.38 | 0.00 | 0.09 |
| W9 | ~food*sex + puuv + deworm + nematode + repro | 232.61 | 0.22 | 0.08 |
| W0 | ~1 | 232.68 | 0.29 | 0.08 |
| W20 | ~puuv*sex + nematode*food + deworm + repro | 233.21 | 0.83 | 0.06 |
| W30 | ~puuv*food*sex + deworm + nematode + repro | 233.31 | 0.93 | 0.06 |
| W24 | ~food*sex + puuv*nematode + deworm + repro | 233.57 | 1.19 | 0.05 |
| W18 | ~puuv*sex + deworm*nematode + food + repro | 233.86 | 1.48 | 0.05 |
| W43 | ~food*deworm*nematode + puuv*sex + repro | 234.19 | 1.81 | 0.04 |
| W21 | ~deworm*nematode + food*sex + puuv + repro | 234.42 | 2.04 | 0.03 |
| W16 | ~puuv*deworm + food*sex + nematode + repro | 234.52 | 2.14 | 0.03 |
| W1 | ~puuv + food + deworm + nematode + sex + repro | 234.80 | 2.42 | 0.03 |
| W10 | ~deworm*sex + puuv + food + nematode + repro | 234.95 | 2.57 | 0.03 |
| W40 | ~puuv*food*sex + deworm*nematode + repro | 234.99 | 2.61 | 0.03 |
| W7 | ~food*deworm + puuv + nematode + sex + repro | 235.11 | 2.73 | 0.02 |
| W23 | ~nematode*food + deworm*sex + puuv + repro | 235.49 | 3.11 | 0.02 |
| W8 | ~nematode*food + puuv + deworm + sex + repro | 235.51 | 3.13 | 0.02 |
| W34 | ~food*deworm*sex + puuv + nematode + repro | 235.72 | 3.34 | 0.02 |
| W39 | ~puuv*deworm*nematode + food*sex + repro | 235.79 | 3.41 | 0.02 |
| W25 | ~deworm*sex + puuv*nematode + food + repro | 235.86 | 3.48 | 0.02 |
| W4 | ~puuv*nematode + food + deworm + sex + repro | 235.88 | 3.50 | 0.02 |
| W17 | ~puuv*nematode + food*deworm + sex + repro | 236.06 | 3.68 | 0.02 |
| W11 | ~nematode*sex + puuv + food + deworm + repro | 236.10 | 3.72 | 0.01 |
| W36 | ~food*nematode*sex + puuv + deworm + repro | 236.18 | 3.80 | 0.01 |
| W44 | ~food*deworm*sex + puuv*nematode + repro | 236.28 | 3.90 | 0.01 |
| W6 | ~deworm*nematode + puuv + food + sex + repro | 236.42 | 4.04 | 0.01 |
| W22 | ~food*deworm + nematode*sex + puuv + repro | 236.53 | 4.15 | 0.01 |
| W3 | ~puuv*deworm + food + nematode + sex + repro | 236.74 | 4.36 | 0.01 |
| W2 | ~puuv*food + deworm + nematode + sex + repro | 236.77 | 4.39 | 0.01 |
| W31 | ~puuv*deworm*sex + food + nematode + repro | 236.90 | 4.52 | 0.01 |
| W33 | ~food*deworm*nematode + puuv + sex + repro | 236.93 | 4.55 | 0.01 |
| W13 | ~puuv*food + deworm*sex + nematode + repro | 236.95 | 4.56 | 0.01 |
| W32 | ~puuv*nematode*sex + food + deworm + repro | 237.18 | 4.80 | 0.01 |
| W42 | ~puuv*nematode*sex + food*deworm + repro | 237.38 | 5.00 | 0.01 |
| W15 | ~puuv*deworm + nematode*food + sex + repro | 237.50 | 5.12 | 0.01 |
| W41 | ~puuv*deworm*sex + food*nematode + repro | 237.65 | 5.27 | 0.01 |
| W38 | ~puuv*food*nematode + deworm*sex + repro | 238.01 | 5.63 | 0.01 |
| W26 | ~nematode*sex + puuv*deworm + food + repro | 238.02 | 5.64 | 0.01 |
| W46 | ~food*nematode*sex + puuv*deworm + repro | 238.11 | 5.73 | 0.01 |
| W14 | ~puuv*food + nematode*sex + deworm + repro | 238.11 | 5.73 | 0.01 |
| W29 | ~puuv*deworm*nematode + food + sex + repro | 238.34 | 5.96 | 0.00 |
| W12 | ~puuv*food + deworm*nematode + sex + repro | 238.42 | 6.04 | 0.00 |
| W35 | ~deworm*nematode*sex + puuv + food + repro | 239.90 | 7.52 | 0.00 |
| W27 | ~puuv*food*deworm + nematode + sex + repro | 240.46 | 8.08 | 0.00 |
| W45 | ~deworm*nematode*sex + puuv*food + repro | 241.95 | 9.57 | 0.00 |

Reproductive status and mortality hazard

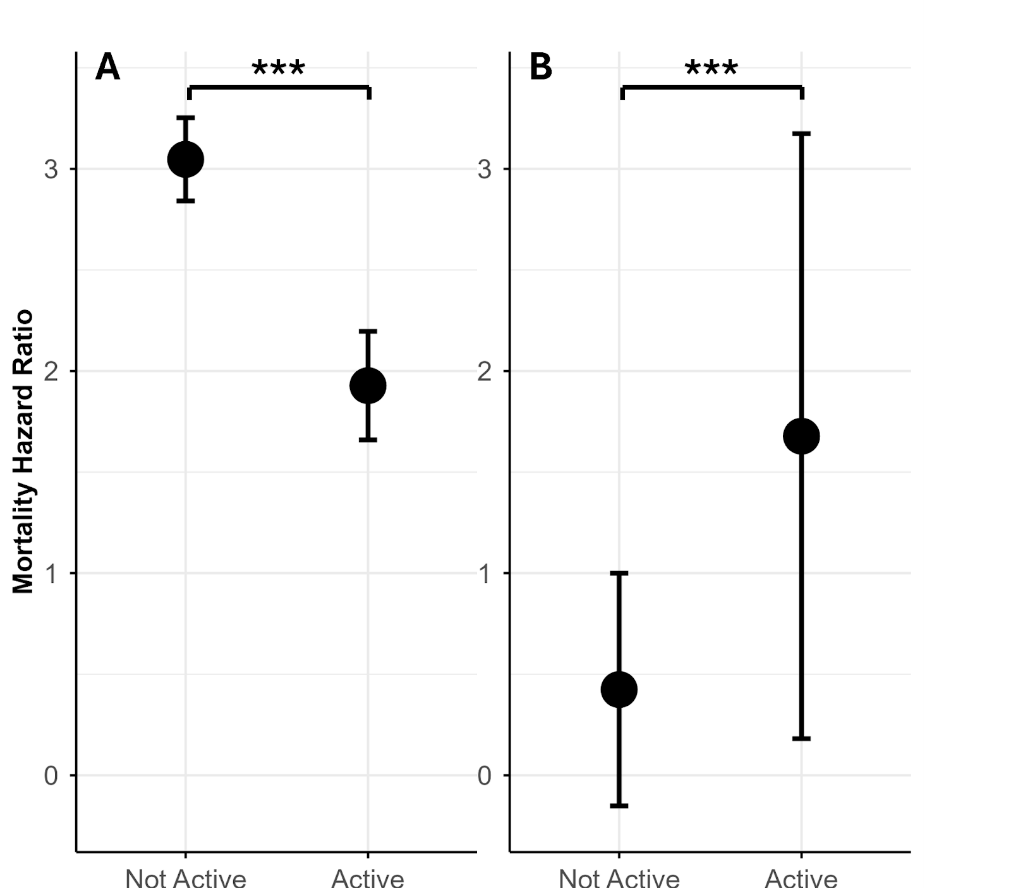

**Supplemental Figure 7:** This figure shows the relationship between reproductive status and relative mortality hazard ratio shown in the primary analysis. The relationship with reproductive status and mortality hazard changed over time (violated the proportional hazards assumption). The direction of the relationship changed when voles were 4.3 months old. This plot shows predicted mortality hazard ratios, relative to the sample average for all predictors in the Cox proportional hazards model (M1). Here Panel A shows the relationship for voles 4 months or younger, while Panel B shows the relationship for voles older than 4 months. Here, M1 was rerun with the data subset into these categories in order to plot the time-varying relationship.

Primarily analysis with transient adult voles removed:

**Supplemental Table 13**: AICc results from running the primary analysis with transient adult voles (reproductively active voles which were only caught once) removed from the dataset. This model comparison was with the set of Cox proportional hazards models (Supplemental Table 5) from the primary analysis run without transient adult voles. Every model listed included a term clustering by individual vole ID, allowing risk error calculations to account for multiple sampling of the same individuals. The parameters “food” and “deworm” refer to the experimental treatment groups, “puuv” refers to PUUV infection status, “nematode” refers to nematode infection status, and “repro” refers to reproductive status. Any term with “tt()” indicates that this term was allowed to vary with time based on the equation tt=x*time (vole age). These time-varying terms were identified by checking the proportional hazards assumption. The first three models were favored equally (ΔAICc <2). T7 had the highest AICc weight (0.18 AICc weight), so it was used for further inference.

| **Model Number** | **Model Predictors** | **AICc** | Δ**AICc** | **AICc Wt** |
| --- | --- | --- | --- | --- |
| T7 | ~food*deworm + puuv + nematode + sex + repro + tt(repro) + month | 5476.35 | 0.00 | 0.18 |
| T17 | ~puuv*nematode + food*deworm + sex + repro + tt(repro) + month | 5477.99 | 1.64 | 0.08 |
| T19 | ~puuv*sex + food*deworm + nematode + repro + tt(repro) + month | 5478.34 | 1.98 | 0.07 |
| T22 | ~food*deworm + nematode*sex + puuv + repro + tt(repro) + month | 5478.37 | 2.01 | 0.07 |
| T1 | ~puuv + food + deworm + nematode + sex + repro + tt(repro) + month | 5478.56 | 2.20 | 0.06 |
| T34 | ~food*deworm*sex + puuv + nematode + repro + tt(repro) + month | 5478.78 | 2.43 | 0.05 |
| T27 | ~puuv*food*deworm + nematode + sex + repro + tt(repro) + month | 5479.42 | 3.07 | 0.04 |
| T3 | ~puuv*deworm + food + nematode + sex + repro + tt(repro) + month | 5479.93 | 3.57 | 0.03 |
| T4 | ~puuv*nematode + food + deworm + sex + repro + tt(repro) + month | 5480.10 | 3.75 | 0.03 |
| T10 | ~deworm*sex + puuv + food + nematode + repro + tt(repro) + month | 5480.15 | 3.80 | 0.03 |
| T6 | ~deworm*nematode + puuv + food + sex + repro + tt(repro) + month | 5480.21 | 3.86 | 0.03 |
| T44 | ~food*deworm*sex + puuv*nematode + repro + tt(repro) + month | 5480.29 | 3.93 | 0.03 |
| T9 | ~food*sex + puuv + deworm + nematode + repro + tt(repro) + month | 5480.41 | 4.06 | 0.02 |
| T8 | ~nematode*food + puuv + deworm + sex + repro + tt(repro) + month | 5480.47 | 4.12 | 0.02 |
| T5 | ~puuv*sex + deworm + nematode + food + repro + tt(repro) + month | 5480.54 | 4.19 | 0.02 |
| T11 | ~nematode*sex + puuv + food + deworm + repro + tt(repro) + month | 5480.55 | 4.20 | 0.02 |
| T2 | ~puuv*food + deworm + nematode + sex + repro + tt(repro) + month | 5480.57 | 4.22 | 0.02 |
| T37 | ~puuv*food*deworm + nematode*sex + repro + tt(repro) + month | 5481.41 | 5.05 | 0.01 |
| T25 | ~deworm*sex + puuv*nematode + food + repro + tt(repro) + month | 5481.65 | 5.29 | 0.01 |
| T33 | ~food*deworm*nematode + puuv + sex + repro + tt(repro) + month | 5481.65 | 5.29 | 0.01 |
| T16 | ~puuv*deworm + food*sex + nematode + repro + tt(repro) + month | 5481.80 | 5.45 | 0.01 |
| T15 | ~puuv*deworm + nematode*food + sex + repro + tt(repro) + month | 5481.83 | 5.47 | 0.01 |
| T26 | ~nematode*sex + puuv*deworm + food + repro + tt(repro) + month | 5481.90 | 5.55 | 0.01 |
| T24 | ~food*sex + puuv*nematode + deworm + repro + tt(repro) + month | 5481.95 | 5.60 | 0.01 |
| T23 | ~nematode*food + deworm*sex + puuv + repro + tt(repro) + month | 5482.06 | 5.70 | 0.01 |
| T21 | ~deworm*nematode + food*sex + puuv + repro + tt(repro) + month | 5482.09 | 5.74 | 0.01 |
| T13 | ~puuv*food + deworm*sex + nematode + repro + tt(repro) + month | 5482.17 | 5.82 | 0.01 |
| T18 | ~puuv*sex + deworm*nematode + food + repro + tt(repro) + month | 5482.21 | 5.86 | 0.01 |
| T12 | ~puuv*food + deworm*nematode + sex + repro + tt(repro) + month | 5482.23 | 5.88 | 0.01 |
| T20 | ~puuv*sex + nematode*food + deworm + repro + tt(repro) + month | 5482.46 | 6.11 | 0.01 |
| T14 | ~puuv*food + nematode*sex + deworm + repro + tt(repro) + month | 5482.57 | 6.21 | 0.01 |
| T43 | ~food*deworm*nematode + puuv*sex + repro + tt(repro) + month | 5483.64 | 7.29 | 0.00 |
| T42 | ~puuv*nematode*sex + food*deworm + repro + tt(repro) + month | 5484.01 | 7.66 | 0.00 |
| T30 | ~puuv*food*sex + deworm + nematode + repro + tt(repro) + month | 5484.09 | 7.73 | 0.00 |
| T31 | ~puuv*deworm*sex + food + nematode + repro + tt(repro) + month | 5484.89 | 8.54 | 0.00 |
| T29 | ~puuv*deworm*nematode + food + sex + repro + tt(repro) + month | 5485.14 | 8.79 | 0.00 |
| T28 | ~puuv*food*nematode + deworm + sex + repro + tt(repro) + month | 5485.60 | 9.25 | 0.00 |
| T40 | ~puuv*food*sex + deworm*nematode + repro + tt(repro) + month | 5485.69 | 9.33 | 0.00 |
| T35 | ~deworm*nematode*sex + puuv + food + repro + tt(repro) + month | 5485.78 | 9.43 | 0.00 |
| T36 | ~food*nematode*sex + puuv + deworm + repro + tt(repro) + month | 5485.99 | 9.63 | 0.00 |
| T32 | ~puuv*nematode*sex + food + deworm + repro + tt(repro) + month | 5486.07 | 9.72 | 0.00 |
| T41 | ~puuv*deworm*sex + food*nematode + repro + tt(repro) + month | 5486.75 | 10.40 | 0.00 |
| T39 | ~puuv*deworm*nematode + food*sex + repro + tt(repro) + month | 5487.04 | 10.69 | 0.00 |
| T38 | ~puuv*food*nematode + deworm*sex + repro + tt(repro) + month | 5487.17 | 10.81 | 0.00 |
| T46 | ~food*nematode*sex + puuv*deworm + repro + tt(repro) + month | 5487.30 | 10.95 | 0.00 |
| T45 | ~deworm*nematode*sex + puuv*food + repro + tt(repro) + month | 5487.80 | 11.45 | 0.00 |
| T0 | ~1 | 5713.49 | 237.13 | 0.00 |

**Supplemental Table 14:** Inference for T7, the model favored in the primary analysis with transient adult voles (reproductively active voles which were only caught once) removed. The results from this model provide information about mortality hazards. The predictors in T7 were “~food*deworm + puuv + nematode + sex + repro + tt(repro) + month”, and the model was clustered by individual vole ID. In these results, the coefficient gives the direction of the relationship between the explanatory variable and the relative hazard of dying (positive indicates increased hazard). The exponentiated coefficient is the hazard ratio and can be used to find the percentage change where 1 is a 0% change in hazard, numbers above 1 are increased hazard (e.g. exp(coef)=1.51 has a 51% higher hazard), and numbers below 1 have decreased hazard (e.g. 0.92 has an 8% decrease in hazard). In T7, the relationship with reproductive status changed direction when voles were 7.7 months old. Refer to Figure 3 for interpretation of the significant Deworm*Fed interaction.

| **Explanatory variable** | **Coefficient** | **Hazard ratio (exponentiated**  **coefficient)** | **Robust**  **standard error** | **P-value** |
| --- | --- | --- | --- | --- |
| PUUV + | 0.53 | 1.70 | 0.10 | 5.9e-8*** |
| Fed | -0.03 | 0.97 | 0.11 | 0.81 |
| Deworm | -0.41 | 0.66 | 0.13 | 0.0020** |
| Nematode+ | -0.30 | 0.74 | 0.10 | 0.0023** |
| Male | 0.16 | 1.17 | 0.08 | 0.063 |
| Repro active | -2.62 | 0.07 | 0.40 | 6.0e-11*** |
| Time varying repro active | 0.34 | 1.40 | 0.08 | 6.1e-5*** |
| Month | -0.47 | 0.62 | 0.04 | <2e-16*** |
| Deworm*Fed | 0.38 | 1.46 | 0.17 | 0.03* |

Secondary analysis with transient adult voles removed:

**Supplemental Table 15**: AICc results from running the secondary analysis with transient adult voles (reproductively active voles which were only caught once) removed from the dataset. This comparison of Cox proportional hazards models (Supplemental Table 5) was run with data from younger voles (<7 months old). Every model listed included a term clustering by individual vole ID, allowing risk error calculations to account for multiple sampling of the same individuals. The parameters “food” and “deworm” refer to the experimental treatment groups, “puuv” refers to PUUV infection status, “nematode” refers to nematode infection status, and “repro” refers to reproductive status. Any term with “tt()” indicates that this term was allowed to vary with time based on the equation tt=x*time (vole age). These time-varying terms were identified by checking the proportional hazards assumption. The first four models were favored equally (ΔAICc <2). TY34 had the highest AICc weight (0.17 AICc weight), so it was used for further inference.

| **Model Number** | **Model Predictors** | **AICc** | Δ**AICc** | **AICc Wt** |
| --- | --- | --- | --- | --- |
| TY34 | ~food*deworm*sex + puuv + nematode + repro + tt(repro) + month | 4465.69 | 0.00 | 0.17 |
| TY7 | ~food*deworm + puuv + nematode + sex + repro + tt(repro) + month | 4466.77 | 1.07 | 0.10 |
| TY44 | ~food*deworm*sex + puuv*nematode + repro + tt(repro) + month | 4467.04 | 1.35 | 0.09 |
| TY19 | ~puuv*sex + food*deworm + nematode + repro + tt(repro) + month | 4467.65 | 1.96 | 0.06 |
| TY1 | ~puuv + food + deworm + nematode + sex + repro + tt(repro) + month | 4467.96 | 2.26 | 0.05 |
| TY17 | ~puuv*nematode + food*deworm + sex + repro + tt(repro) + month | 4468.27 | 2.57 | 0.05 |
| TY22 | ~food*deworm + nematode*sex + puuv + repro + tt(repro) + month | 4468.38 | 2.69 | 0.04 |
| TY5 | ~puuv*sex + deworm + nematode + food + repro + tt(repro) + month | 4468.85 | 3.15 | 0.04 |
| TY9 | ~food*sex + puuv + deworm + nematode + repro + tt(repro) + month | 4469.25 | 3.56 | 0.03 |
| TY10 | ~deworm*sex + puuv + food + nematode + repro + tt(repro) + month | 4469.30 | 3.60 | 0.03 |
| TY4 | ~puuv*nematode + food + deworm + sex + repro + tt(repro) + month | 4469.48 | 3.78 | 0.03 |
| TY11 | ~nematode*sex + puuv + food + deworm + repro + tt(repro) + month | 4469.54 | 3.84 | 0.02 |
| TY3 | ~puuv*deworm + food + nematode + sex + repro + tt(repro) + month | 4469.67 | 3.98 | 0.02 |
| TY2 | ~puuv*food + deworm + nematode + sex + repro + tt(repro) + month | 4469.69 | 4.00 | 0.02 |
| TY8 | ~nematode*food + puuv + deworm + sex + repro + tt(repro) + month | 4469.88 | 4.19 | 0.02 |
| TY6 | ~deworm*nematode + puuv + food + sex + repro + tt(repro) + month | 4469.95 | 4.25 | 0.02 |
| TY27 | ~puuv*food*deworm + nematode + sex + repro + tt(repro) + month | 4470.07 | 4.37 | 0.02 |
| TY24 | ~food*sex + puuv*nematode + deworm + repro + tt(repro) + month | 4470.75 | 5.06 | 0.01 |
| TY20 | ~puuv*sex + nematode*food + deworm + repro + tt(repro) + month | 4470.77 | 5.08 | 0.01 |
| TY25 | ~deworm*sex + puuv*nematode + food + repro + tt(repro) + month | 4470.83 | 5.14 | 0.01 |
| TY18 | ~puuv*sex + deworm*nematode + food + repro + tt(repro) + month | 4470.84 | 5.15 | 0.01 |
| TY16 | ~puuv*deworm + food*sex + nematode + repro + tt(repro) + month | 4470.98 | 5.29 | 0.01 |
| TY13 | ~puuv*food + deworm*sex + nematode + repro + tt(repro) + month | 4471.03 | 5.33 | 0.01 |
| TY23 | ~nematode*food + deworm*sex + puuv + repro + tt(repro) + month | 4471.22 | 5.53 | 0.01 |
| TY21 | ~deworm*nematode + food*sex + puuv + repro + tt(repro) + month | 4471.24 | 5.55 | 0.01 |
| TY26 | ~nematode*sex + puuv*deworm + food + repro + tt(repro) + month | 4471.27 | 5.57 | 0.01 |
| TY14 | ~puuv*food + nematode*sex + deworm + repro + tt(repro) + month | 4471.29 | 5.59 | 0.01 |
| TY15 | ~puuv*deworm + nematode*food + sex + repro + tt(repro) + month | 4471.60 | 5.90 | 0.01 |
| TY37 | ~puuv*food*deworm + nematode*sex + repro + tt(repro) + month | 4471.64 | 5.94 | 0.01 |
| TY12 | ~puuv*food + deworm*nematode + sex + repro + tt(repro) + month | 4471.67 | 5.98 | 0.01 |
| TY42 | ~puuv*nematode*sex + food*deworm + repro + tt(repro) + month | 4472.24 | 6.55 | 0.01 |
| TY33 | ~food*deworm*nematode + puuv + sex + repro + tt(repro) + month | 4472.52 | 6.83 | 0.01 |
| TY32 | ~puuv*nematode*sex + food + deworm + repro + tt(repro) + month | 4473.31 | 7.62 | 0.00 |
| TY30 | ~puuv*food*sex + deworm + nematode + repro + tt(repro) + month | 4473.42 | 7.72 | 0.00 |
| TY43 | ~food*deworm*nematode + puuv*sex + repro + tt(repro) + month | 4473.42 | 7.73 | 0.00 |
| TY36 | ~food*nematode*sex + puuv + deworm + repro + tt(repro) + month | 4473.65 | 7.96 | 0.00 |
| TY31 | ~puuv*deworm*sex + food + nematode + repro + tt(repro) + month | 4473.91 | 8.22 | 0.00 |
| TY28 | ~puuv*food*nematode + deworm + sex + repro + tt(repro) + month | 4474.37 | 8.68 | 0.00 |
| TY29 | ~puuv*deworm*nematode + food + sex + repro + tt(repro) + month | 4474.39 | 8.70 | 0.00 |
| TY35 | ~deworm*nematode*sex + puuv + food + repro + tt(repro) + month | 4474.75 | 9.05 | 0.00 |
| TY46 | ~food*nematode*sex + puuv*deworm + repro + tt(repro) + month | 4475.37 | 9.67 | 0.00 |
| TY40 | ~puuv*food*sex + deworm*nematode + repro + tt(repro) + month | 4475.42 | 9.73 | 0.00 |
| TY39 | ~puuv*deworm*nematode + food*sex + repro + tt(repro) + month | 4475.76 | 10.06 | 0.00 |
| TY38 | ~puuv*food*nematode + deworm*sex + repro + tt(repro) + month | 4475.78 | 10.08 | 0.00 |
| TY41 | ~puuv*deworm*sex + food*nematode + repro + tt(repro) + month | 4475.83 | 10.14 | 0.00 |
| TY45 | ~deworm*nematode*sex + puuv*food + repro + tt(repro) + month | 4476.47 | 10.78 | 0.00 |
| TY0 | ~1 | 4725.26 | 259.57 | 0.00 |

**Supplemental Table 16**: AICc results from running the secondary analysis with transient adult voles (reproductively active voles which were only caught once) removed from the dataset. This comparison of Cox proportional hazards models (Supplemental Table 5) was run with data from older voles (7 months or older). Every model listed included a term clustering by individual vole ID, allowing risk error calculations to account for multiple sampling of the same individuals. The parameters “food” and “deworm” refer to the experimental treatment groups, “puuv” refers to PUUV infection status, “nematode” refers to nematode infection status, and “repro” refers to reproductive status. None of the relationships with predictors varied with time. The first seven models were favored equally (ΔAICc <2) and were not within 2 units of the null model. TO9 had the highest AICc weight (0.11 AICc weight), so it was used for further inference.

| **Model Number** | **Model Predictors** | **AICc** | Δ**AICc** | **AICc Wt** |
| --- | --- | --- | --- | --- |
| TO9 | ~food*sex + puuv + deworm + nematode + repro + month | 930.47 | 0.00 | 0.11 |
| TO1 | ~puuv + food + deworm + nematode + sex + repro + month | 930.74 | 0.27 | 0.10 |
| TO16 | ~puuv*deworm + food*sex + nematode + repro + month | 931.72 | 1.25 | 0.06 |
| TO3 | ~puuv*deworm + food + nematode + sex + repro + month | 932.09 | 1.62 | 0.05 |
| TO5 | ~puuv*sex + deworm + nematode + food + repro + month | 932.13 | 1.66 | 0.05 |
| TO24 | ~food*sex + puuv*nematode + deworm + repro + month | 932.26 | 1.79 | 0.05 |
| TO21 | ~deworm*nematode + food*sex + puuv + repro + month | 932.41 | 1.94 | 0.04 |
| TO4 | ~puuv*nematode + food + deworm + sex + repro + month | 932.59 | 2.12 | 0.04 |
| TO6 | ~deworm*nematode + puuv + food + sex + repro + month | 932.64 | 2.17 | 0.04 |
| TO8 | ~nematode*food + puuv + deworm + sex + repro + month | 932.65 | 2.18 | 0.04 |
| TO10 | ~deworm*sex + puuv + food + nematode + repro + month | 932.73 | 2.26 | 0.04 |
| TO7 | ~food*deworm + puuv + nematode + sex + repro + month | 932.81 | 2.34 | 0.04 |
| TO11 | ~nematode*sex + puuv + food + deworm + repro + month | 932.82 | 2.35 | 0.04 |
| TO2 | ~puuv*food + deworm + nematode + sex + repro + month | 932.83 | 2.36 | 0.03 |
| TO31 | ~puuv*deworm*sex + food + nematode + repro + month | 933.81 | 3.34 | 0.02 |
| TO18 | ~puuv*sex + deworm*nematode + food + repro + month | 933.98 | 3.51 | 0.02 |
| TO20 | ~puuv*sex + nematode*food + deworm + repro + month | 934.04 | 3.57 | 0.02 |
| TO15 | ~puuv*deworm + nematode*food + sex + repro + month | 934.05 | 3.58 | 0.02 |
| TO26 | ~nematode*sex + puuv*deworm + food + repro + month | 934.19 | 3.71 | 0.02 |
| TO19 | ~puuv*sex + food*deworm + nematode + repro + month | 934.21 | 3.74 | 0.02 |
| TO25 | ~deworm*sex + puuv*nematode + food + repro + month | 934.62 | 4.15 | 0.01 |
| TO23 | ~nematode*food + deworm*sex + puuv + repro + month | 934.64 | 4.17 | 0.01 |
| TO17 | ~puuv*nematode + food*deworm + sex + repro + month | 934.67 | 4.20 | 0.01 |
| TO12 | ~puuv*food + deworm*nematode + sex + repro + month | 934.74 | 4.26 | 0.01 |
| TO13 | ~puuv*food + deworm*sex + nematode + repro + month | 934.83 | 4.35 | 0.01 |
| TO22 | ~food*deworm + nematode*sex + puuv + repro + month | 934.91 | 4.44 | 0.01 |
| TO14 | ~puuv*food + nematode*sex + deworm + repro + month | 934.92 | 4.45 | 0.01 |
| TO0 | ~1 | 935.47 | 5.00 | 0.01 |
| TO30 | ~puuv*food*sex + deworm + nematode + repro + month | 935.70 | 5.23 | 0.01 |
| TO41 | ~puuv*deworm*sex + food*nematode + repro + month | 935.73 | 5.25 | 0.01 |
| TO34 | ~food*deworm*sex + puuv + nematode + repro + month | 935.99 | 5.51 | 0.01 |
| TO36 | ~food*nematode*sex + puuv + deworm + repro + month | 936.29 | 5.81 | 0.01 |
| TO40 | ~puuv*food*sex + deworm*nematode + repro + month | 937.58 | 7.11 | 0.00 |
| TO33 | ~food*deworm*nematode + puuv + sex + repro + month | 937.62 | 7.15 | 0.00 |
| TO39 | ~puuv*deworm*nematode + food*sex + repro + month | 937.62 | 7.15 | 0.00 |
| TO46 | ~food*nematode*sex + puuv*deworm + repro + month | 937.64 | 7.16 | 0.00 |
| TO44 | ~food*deworm*sex + puuv*nematode + repro + month | 937.82 | 7.35 | 0.00 |
| TO29 | ~puuv*deworm*nematode + food + sex + repro + month | 937.99 | 7.52 | 0.00 |
| TO32 | ~puuv*nematode*sex + food + deworm + repro + month | 938.07 | 7.59 | 0.00 |
| TO27 | ~puuv*food*deworm + nematode + sex + repro + month | 938.25 | 7.78 | 0.00 |
| TO28 | ~puuv*food*nematode + deworm + sex + repro + month | 938.31 | 7.83 | 0.00 |
| TO35 | ~deworm*nematode*sex + puuv + food + repro + month | 938.64 | 8.16 | 0.00 |
| TO43 | ~food*deworm*nematode + puuv*sex + repro + month | 939.13 | 8.66 | 0.00 |
| TO42 | ~puuv*nematode*sex + food*deworm + repro + month | 940.19 | 9.71 | 0.00 |
| TO38 | ~puuv*food*nematode + deworm*sex + repro + month | 940.32 | 9.85 | 0.00 |
| TO37 | ~puuv*food*deworm + nematode*sex + repro + month | 940.39 | 9.91 | 0.00 |
| TO45 | ~deworm*nematode*sex + puuv*food + repro + month | 940.77 | 10.30 | 0.00 |

**Supplemental Table 17:** Inference for the younger and older vole analyses with transient adult voles (reproductively active voles which were only caught once) removed. Candidate models run in the primary analysis were run for younger voles (less than 7 months) or older voles (7 months or older) with transient adult voles removed. The results from these models provide information about mortality hazards. For younger voles, TY34 (model predictors: “~food*deworm*sex + puuv + nematode + repro + tt(repro) + month”) was favored with the highest AICc weight. For older voles, TO9 (model predictors: “~food*sex + puuv + deworm + nematode + repro + month”) was favored with the highest AICc weight. In these results, the coefficient gives the direction of the relationship between the explanatory variable and the relative hazard of dying (positive indicates increased hazard). The exponentiated coefficient is the hazard ratio and can be used to find the percentage change where 1 is a 0% change in hazard, numbers above 1 are increased hazard (e.g. exp(coef)=1.51 has a 51% higher hazard), and numbers below 1 have decreased hazard (e.g. 0.92 has an 8% decrease in hazard). In TY34, the relationship with reproductive status changed direction when voles were 6.2 months old. See Supplemental Figure 8 for interpretation of the Deworm*Male*Fed interaction in TY34. See Supplemental Figure 9 for interpretation of the Male*Fed interaction in TO9.

**Younger voles:**

| **Explanatory variable** | **Coefficient** | **Hazard ratio (exponentiated coefficient)** | **Robust**  **standard**  **error** | **P-value** |
| --- | --- | --- | --- | --- |
| PUUV + | 0.55 | 1.73 | 0.11 | 1.6e-6*** |
| Fed | 0.18 | 1.20 | 0.21 | 0.38 |
| Deworm | -0.01 | 0.99 | 0.26 | 0.98 |
| Nematode+ | -0.25 | 0.78 | 0.11 | 0.026* |
| Male | 0.28 | 1.32 | 0.20 | 0.16 |
| Repro active | -4.55 | 0.01 | 0.60 | 4.2e-14*** |
| Time varying repro active | 0.74 | 2.09 | 0.13 | 2.6e-8*** |
| Month | -0.60 | 0.55 | 0.04 | <2e-16*** |
| Deworm*Fed | -0.16 | 0.85 | 0.31 | 0.60 |
| Male*Fed | -0.18 | 0.84 | 0.26 | 0.49 |
| Deworm*Male | -0.88 | 0.41 | 0.34 | 0.010* |
| Deworm*Male*Fed | 1.03 | 2.79 | 0.41 | 0.013* |

**Older voles:**

| **Explanatory variable** | **Coefficient** | **Hazard ratio (exponentiated coefficient)** | **Robust**  **standard**  **error** | **P-value** |
| --- | --- | --- | --- | --- |
| PUUV + | -0.03 | 0.97 | 0.18 | 0.85 |
| Fed | -0.65 | 0.52 | 0.28 | 0.019* |
| Deworm | 0.03 | 1.03 | 0.17 | 0.87 |
| Nematode+ | -0.33 | 0.72 | 0.19 | 0.08 |
| Male | 0.17 | 1.19 | 0.24 | 0.46 |
| Repro active | 1.10 | 3.01 | 0.43 | 0.010** |
| Month | 0.13 | 1.14 | 0.10 | 0.18 |
| Male*Fed | 0.60 | 1.82 | 0.35 | 0.089 |

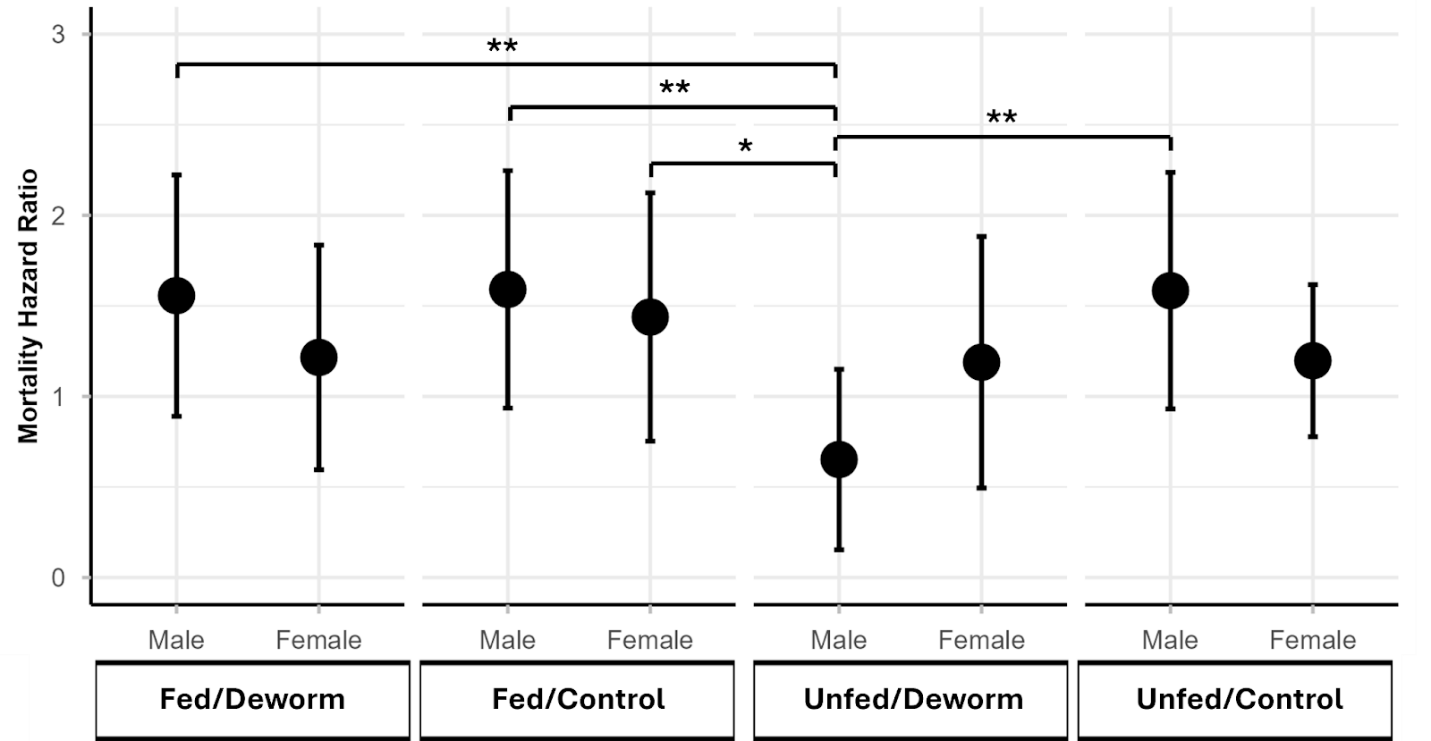

**Supplemental Figure 8:** Plot of significant interaction between food treatment, deworm treatment and sex in TY34 (the top favored models with the highest AICc weight in the younger vole analysis with transient voles removed). This plot shows predicted mortality hazard ratios, relative to the sample average for all predictors in the Cox proportional hazards model (TY34). Among male voles, we detected that receiving deworm reduces the relative mortality hazard, but only when food is not added to the environment. Significance was calculated from post hoc pairwise comparisons (with Bonferroni correction).

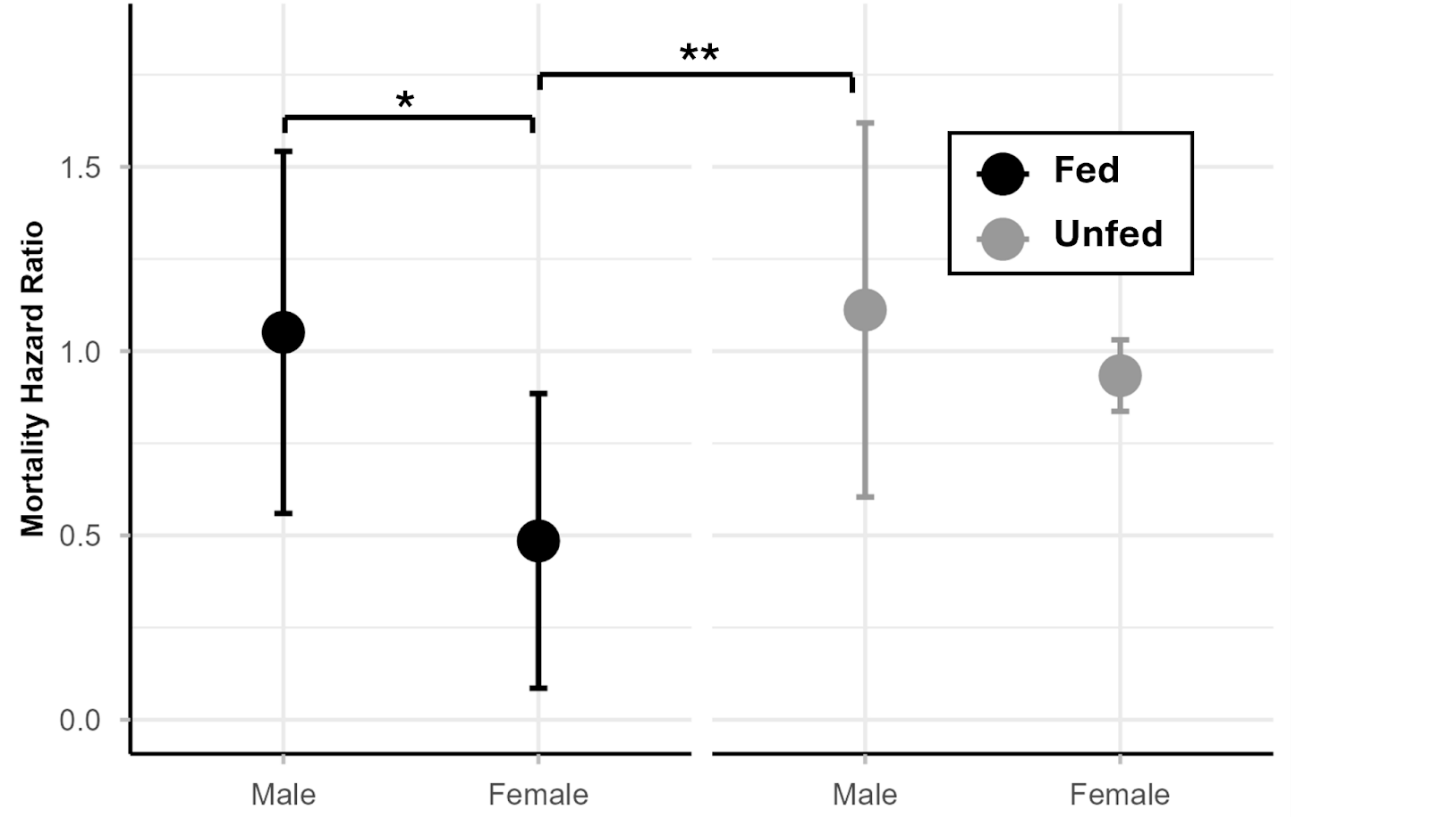

**Supplemental Figure 9:** Plot of nonsignificant interaction between food treatment and sex in TO9 (the top favored model with the highest AICc weight in the older vole analysis with transient voles removed). This plot shows predicted mortality hazard ratios, relative to the sample average for all predictors in the Cox proportional hazards model (TO9). We only detected a significant difference in relative mortality hazard between male and female voles at fed sites. Significance was calculated from post hoc pairwise comparisons (with Bonferroni correction).

**Supplementary Materials References**
